# MAssively-Parallel Flow cytometry Xplorer (MAPFX): A Toolbox for Analysing Data from the Massively-Parallel Cytometry Experiments

**DOI:** 10.1101/2024.02.28.582452

**Authors:** Hsiao-Chi Liao, Terence P. Speed, Davis J. McCarthy, Agus Salim

## Abstract

Massively-Parallel Cytometry (MPC) experiments allow cost-effective quantification of more than 200 surface proteins at single-cell resolution. The Infinity Flow (Inflow) analysis protocol was developed to measure highly informative protein ‘*backbone*’ markers on all cells in all wells distributed across three 96-well plates, along with well-specific exploratory protein ‘*infinity*’ markers. Backbone markers can be used to impute the infinity markers on cells in all other wells using machine learning methods. This protocol offers unprecedented opportunities for more comprehensive classification of cell types. However, some aspects of the protocol can be improved, including methods for background correction and removal of unwanted variation. Here, we propose *MAPFX* as an end-to-end toolbox that carefully pre-processes the raw data from MPC experiments, and further imputes the ‘missing’ infinity markers in the wells without those measurements. Our pipeline starts by performing background correction on raw intensities to remove the noise from electronic baseline restoration and fluorescence compensation by adapting a normal-exponential convolution model. Unwanted technical variation, from sources such as well effects, is then removed using a log-normal model with plate, column, and row factors, after which infinity markers are imputed using the informative backbone markers as predictors. The completed dataset can then be used for clustering and other statistical analyses. Unique features of our approach include performing background correction prior to imputation and removing unwanted variation from the data at the cell-level, while explicitly accounting for the potential association between biology and unwanted factors. We benchmark our pipeline against alternative pipelines and demonstrate that our approach is better at preserving biological signals, removing unwanted variation, and imputing unmeasured infinity markers.

## Introduction

### Technologies for profiling single-cell surface proteins

The heterogeneity of cell-surface proteins helps in identifying cellular subpopulations in the immune system. To date, more than 400 cluster of differentiation (CD) markers have been identified.^1^ Therefore, the ability to simultaneously measure the expression of a large number of such markers at the single-cell level provides an opportunity to discover more refined cell subtypes as well as rare cell types. Modern technologies such as conventional flow cytometry (with fluorescent tags) and mass cytometry (CyTOF, with metal tags) can measure multiple surface proteins simultaneously for each individual cell. However, due to limitations in these technologies, only up to 17 cell parameters can be measured by flow cytometry,^2^ while CyTOF allows up to around 40 cell parameters to be quantified.^3^ Massively-parallel cytometry (MPC) experimental techniques have been developed to enable simultaneous measurements of more surface markers. MPC is a novel form of flow cytometry that can measure hundreds of cell-surface proteins in low cost plate-based antibody screening panels such as the LEGENDScreen PE kit from BioLegend, and allows larger cell throughput than the conventional fluorescence and mass cytometry (CyTOF) based technologies.^4^ In an MPC experiment, certain well-characterised proteins are measured for all cells in every well of the plates (termed ‘*backbone*’), and unique exploratory marker proteins (termed ‘*infinity*’) are measured sparsely across wells. Becht *et al*. (2021) have developed methods for imputing unmeasured exploratory markers by constructing relationships between the backbone markers and the infinity markers within each well to obtain a completed dataset for downstream analyses.^4^

### Computational and statistical methods for analysing single-cell proteomics data from MPC experiments

Most studies have treated MPC data as outputs from multiple conventional flow cytometry,^5,6, 7^ and analyse the data using software such as FlowJo™ (BD Biosciences) to apply gating strategies for identifying cell types. However, Becht *et al*. (2021) not only performed gating on their MPC dataset, but also imputed the unmeasured exploratory proteins for a more comprehensive analysis using the imputed dataset. To obtain a completed dataset with all measurements on all cells, they established the relationship between backbone markers and well-specific infinity markers using linear and non-linear models trained on cells from one well, followed by applying the models to cells from the other wells to impute the unmeasured infinity markers. Non-linear models outperformed the linear model in their experiments, indicating that the relationships between the backbone markers and the infinity markers are non-linear. The non-linear models used were LASSO3 (degree 3 polynomial regression with L1 regularisation), NN (Neural Network), SVM (Support Vector Machine), and XGBoost (eXtreme Gradient Boosting); and they found XGBoost returned more accurate results and had the shortest running time compared to other non-linear models. After imputation, they used the completed dataset to derive cell clusters using the PhenoGraph algorithm^8^ and visualised the result in UMAP coordinates by embedding high-dimensional features.^9^ Importantly, they reported that some subtypes of cells that could not be found using only the backbone features could be identified by analysing the completed data matrix (shown in Fig. 4A of their paper),^4^ demonstrating the value of imputation.

### Removing or reducing unwanted variation from the data from MPC experiments

Pre-processing is an important step for any omics datasets, and unwanted variation due to factors such as batch effects and technological heterogeneity should be reduced as much as possible during this process. Failure to remove unwanted variation may lead to misleading conclusions and false discoveries from downstream analyses. Currently, in the Inflow analysis protocol, Becht *et al*. (2021)^4^ uses the Logicle transformation^10^ followed by a z transformation (zero mean and unit variance) for adjusting the measurements from different wells. The Logicle functions are generalised hyperbolic sine functions (i.e., biexponential functions) with data-driven parameters that give a linear-like transformation for the values around zero, while the positive and negative values far away from zero are transformed to a logarithm-like scale. With this property, the transformed values may be better for the display of cells in scatter plots^11,10^ and thus help in identifying cell populations. However, the inconsistent scales between proteins can result in artifacts when it comes to calculating the distances between cells, which is often required for downstream analyses such as cluster analysis. Although the issue of the inconsistent scales can be fixed by applying the z transformation at protein-level, it forces the mean and variance of every protein marker to be the same, neglecting the fact that marker-specific variation can be exploited to help the imputation.

Since the protein intensities measured from flow cytometry are a mixture of signal and noise, Becht *et al*. (2021) performed background correction on their imputed infinity markers. Their background corrected values are scaled residuals from a regression model that adjusted for the effect measured in the corresponding isotype control well. Isotype controls are antibodies with unknown specificity, and are often used to measure background noise.^12^ However, there can be at least two problems with their operation. First, imputation involves uncertainty. Correcting background noise on the imputed data cannot separate background noise from the noise introduced by imputation, and thus it may over- or under-correct the data. Second, we have found that the effect estimated from the isotype control wells is not always a good measure of the background noise for some infinity markers, as the measurement of noise from the corresponding isotype control well can be larger than that of the measured protein.

To tackle the above issues, we developed *mapfx*.*norm* in our *MAPFX* (MAssively-Parallel Flow cytometry Xplorer) toolbox that adapts the normal-exponential convolution model to remove background noise and to deal with the negative values due to baseline restoration and incomplete fluorescence compensation. Additionally, the well effects from this plate-based experiment are undesirable, and *mapfx*.*norm* uses a log-normal regression model to remove them, while adequately preserving the biological variation in the data. The normalised data is then transformed to the (common) natural logarithm scale for further analysis, resulting in a consistent scale for the data. Finally, *MAPFX* imputes the unmeasured infinity markers and performs cluster analysis using both the adjusted backbone features and the completed dataset. As an end-to-end toolbox for analysing MPC data, *MAPFX* contains the functions for normalisation, imputation, and clustering.

## Methods

Our proposed method, *mapfx*.*norm* was designed for normalising single-cell proteomics data from MPC experiments. Since the backbone markers in MPC experiments can be treated as the protein markers in conventional fluorescence flow cytometry (FFC) experiments, *mapfx*.*norm* can be applied to normalise the protein measurements from FFC assays as well. The data normalisation steps include background correction, removal of unwanted variation, and data transformation. *MAPFX* was implemented as an R package published on GitHub (see “Data Availability” section). We compared our proposed normalisation method (*mapfx*.*norm*) from our *MAPFX* toolbox with two alternatives *lgc*.*z* (Logicle transformation + z, from the *Inflow protocol* ^*4*^) and *lgc*.*comb*.*bio* (Logicle transformation + ComBat^13^). The experimental and the computational pipelines show the details of the Inflow protocol (Fig. 1).

**Figure 1.**
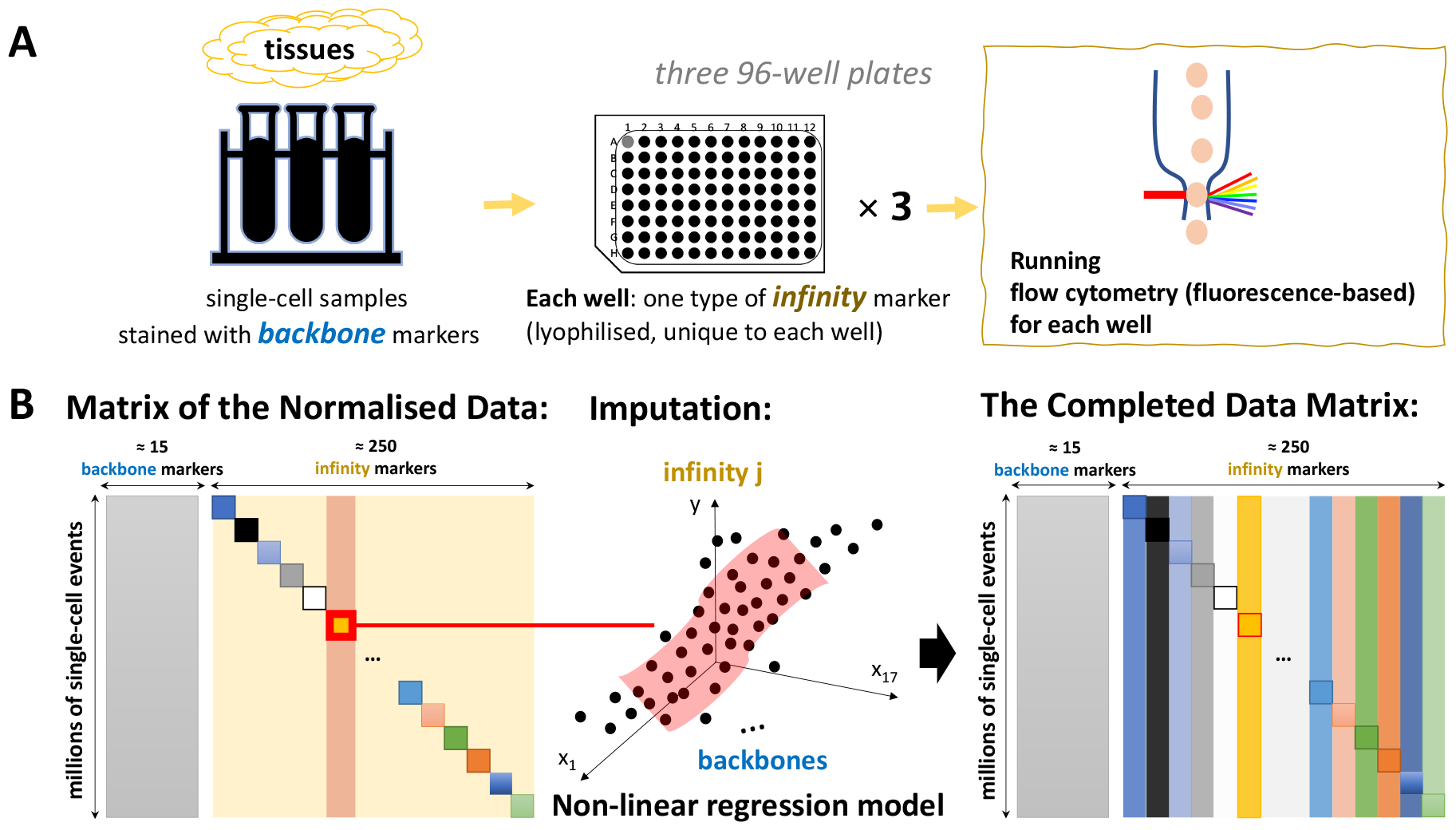
The experimental and the computational pipeline of the Inflow protocol. (A) Experimental pipeline. The single-cell samples are stained with backbone markers (backbone panel staining), then the stained samples are allocated to wells with one particular infinity marker (infinity panel staining), lastly, data can be acquired from the flow cytometry assay for each well. (B) Computational pipeline. The matrix of the normalised data showing that the backbone matrix (gray) contains values for every single-cell (row), but only block diagonal entries of the infinity matrix (yellow) have measurements. Imputation of the unmeasured infinity markers is done by using the backbone markers as predictors in regression models. Finally, the completed data matrix is obtained after imputation.

### The workflow of *mapfx*.*norm* and the alternatives

Our proposed method, *mapfx*.*norm*, cleans up the data before performing imputation because we believe that data should be properly normalised before any downstream statistical procedures are applied. However, the alternative methods *lgc*.*z* and *lgc*.*comb*.*bio* perform the background correction on the imputed data, such that corrected values contain uncertainty introduced during the imputation process. With these strategies, the background-corrected intensities are obtained by regressing out the effect from the corresponding imputed isotype values, whereas the *mapfx*.*norm* workflow normalises the data before undertaking imputation (Supp. Fig. 1). Here, we discuss the key differences between our proposed method (*mapfx*.*norm*) and the alternatives (*lgc*.*z* and *lgc*.*comb*.*bio*) in the following sections on background correction, removal of unwanted variation, and data transformation.

### Background correction

#### mapfx.norm

We adapted the normal-exponential convolution model^14^ for correcting background noise in the measurements of both backbone and infinity markers. With the normal-exponential convolution model, we assume the background noise follows a normal distribution with mean *µ* and variance *σ*^2^, and the true signal follows an exponential distribution with a mean parameter *α*.

The background-corrected value for each protein marker is obtained using the following formula:

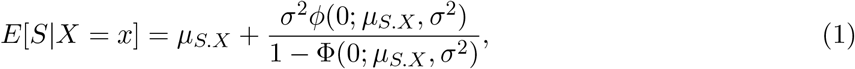

where 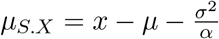, *X* is the random variable that represents the observed protein intensity, and *S* is the random variable that represents its true signal. *ϕ*(.) is the probability density function (PDF) of a normal distribution, and Φ(.) is the cumulative distribution function (CDF) of a normal distribution, each with the specified mean and variance.

The parameters *µ, σ*, and *α* are estimated using Maximum Likelihood Estimation (MLE) using suitable order statistics. Here, we assume the small values of each protein marker are dominated by noise, and the large values are mainly signal. Suppose that we have *N* observations *X* = (*x*_1_, *x*_2_, …, *x*_*N*_). For estimating the parameters of the noise component, we assume that the values *x*_(1)_, *x*_(2)_, …, *x*_(*J*)_ of the first *J* order statistics of *X* are known and contain purely noise, while the remaining *N-J* values are unknown. The likelihood function for the estimation of the noise parameters is written as:

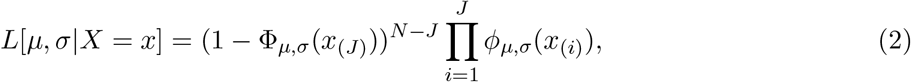

where *ϕ*_*µ,σ*_(.) is the PDF of the normal distribution with the mean *µ* and the standard deviation *σ* and Φ_*µ,σ*_(.) is the CDF of the normal distribution with the mean *µ* and the standard deviation *σ*.

For estimating the parameters of the signal component, we assume that the values of the last *K* order statistics *x*_(*N* −*K*+1)_, *x*_(*N* −*K*+2)_, …, *x*_(*N*)_ of *X* are known and contain purely signal, while the exact signal for the other *N-K* observations are unknown. The likelihood function for the estimation of the signal parameter is then written as:

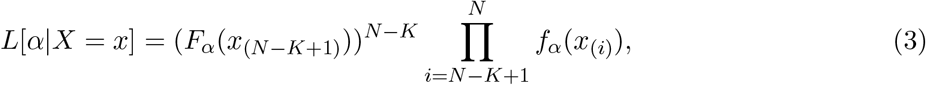

where *f*_*α*_(.) is the PDF of the exponential distribution with the mean parameter *α* and *F*_*α*_(.) is the CDF of the exponential distribution with the mean parameter *α*. In our applications, we found the following settings yielded reasonable results. For estimating noise parameters, we used cells with values smaller than the 10th percentile for each marker, and for estimating parameters of the signal component, we used cells with values larger than the 90th percentile for backbone markers and cells with values larger than 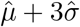 for infinity markers.

#### lgc.z and lgc.comb.bio

Both approaches fit the following regression model to the imputed dataset:

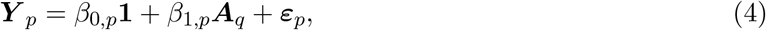

where ***Y*** _*p*_(*N ×* 1) is the raw imputed infinity marker *p* for *N* cells, ***A***_*q*_(*N ×* 1) is infinity marker *p*’s corresponding imputed isotype control, *β*_0,*p*_ is the intercept term, *β*_1,*p*_ is the effect size of the isotype *q* on the infinity marker *p*, and *ε*_*p*_(*N ×* 1) is the error term for the infinity marker *p*.

The background-corrected intensities are then the scaled residuals:

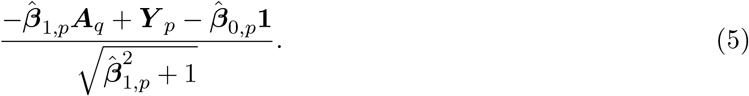

### Removal of unwanted variation

#### mapfx.norm

Here, we assume that the well-effect is the major unwanted factor and that the well-effect can be approximated as the sum of plate, column, and row effects. With the additive model, it only takes 20 (2 for plates, 11 for columns, and 7 for rows) degrees of freedom to estimate the unwanted effects. Note that we fitted another model that considers the interaction between wells by estimating 265 parameters (266 wells - 1) and we found that it did not outperform the simpler model. Therefore, the additive model is adapted to our approach. The log-normal regression model for the adjustment of the well effect is as follows:

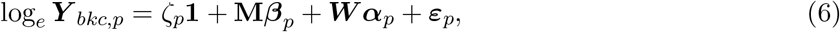

where log ***Y*** _*bkc,p*_(*N ×* 1) is the vector of the background-corrected values of protein *p* for *N* cells, ***M*** (*N × m*) is the design matrix that contains information about (*m* + 1) biological clusters, with ***M*** (*c, j*) = 1 if the *c*th cell is part of the *j*th biological cluster and 0 otherwise, **W**(*N ×* 20) is the design matrix for the known unwanted factors, consisting of two dummy variables for the plate factor, seven for the row factor, and eleven for the column factor, ***β***_*p*_(*m ×* 1) is the vector of biological coefficients with the sum-to-zero constraint 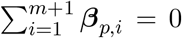, and ***α***_*p*_(20 *×* 1) is the vector of the coefficients for unwanted factors with the sum-to-zero constraint 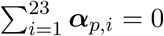, *ζ*_*p*_ is the mean intensity for protein *p*, and finally ***ε***_*p*_(*N ×* 1) is the vector of error terms for protein *p*. The values after adjusting for the well effect can be obtained with the following formula:

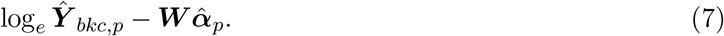

#### lgc.z

For protein *p*, cell *c* in well *w*, the adjusted data is obtained using the following formula:

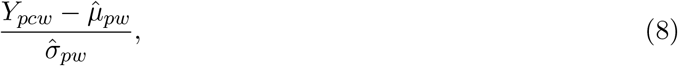

where *Y*_*pcw*_ is the Logicle transformed value, 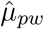 is the marker-specific mean estimated from well *w* for protein *p*, and 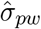 is the marker-specific standard deviation estimated from well *w* for protein *p*.

#### lgc.comb.bio

For protein *p*, cell *c* in well *w*, the adjusted data is obtained using the following formula:

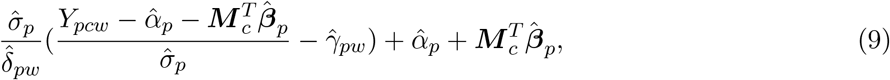

where *Y*_*pcw*_ is the Logicle transformed value, 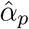 and 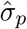 are the estimated mean value and standard deviation of the protein intensity *p*, ***M*** _*c*_(*m ×* 1) is the biological information such as cluster labelling for cell *c* and 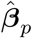 is the corresponding vector of regression coefficients. The 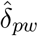 and 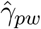 represent multiplicative and additive well effects of well *w* for protein *p*, respectively (see Johnson and Li (2006)^13^ for further details).

### Data transformation

#### **mapfx.norm** uses the following logarithmic transformation

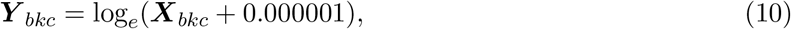

where ***X***_*bkc*_(*N ×* 1) are the non-negative background corrected values for *N* cells, and ***Y*** _*bkc*_(*N ×* 1) are the transformed values on the natural logarithm scale. A tiny offset value 0.000001 is set to not distort the non-negative background corrected values too much.

#### **lgc.z and lgc.comb.bio** use the Logicle transformation

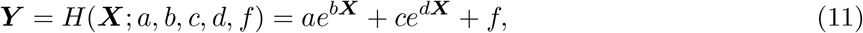

where ***X***(*N ×* 1) is the raw protein intensities for *N* cells, ***Y*** (*N ×* 1) is the values on the Logicle scale, H(.) is a general expression of the Logicle function which is a generalised hyperbolic sine function, and *a, b, c, d, f* are data-driven parameters (see Parks *et al*. (2006)^10^ for further details).

### Regression models and their hyperparameters for imputation

*MAPFX* adapts the models that Becht *et al*. (2021) used for imputing infinity markers, including one linear regression and three non-linear regression models. The parameter settings for the *R* functions are as follows. The *linear* model is fitted by using the *stats::lm* function, the *lasso3* model is fitted using the *glmnetUtils::glmnet* function, with the parameter of *L*_1_-penalty automatically chosen using 10-fold cross-validation. The *svm* model is fitted using the *e1071::svm* function with nu = 0.5 and a radial basis function (RBF) kernel. Finally, the *xgboost* model is fitted using the *xgboost::xgboost* function with `nrounds (maximum number of iterations) = 1500 and eta (the learning rate)= 0.03`. Following Becht *et al*. (2021), for each measured infinity marker, *MAPFX* uses half of the cells to train the model and uses the other half as the testing set. Only cells in the testing sets are retained for the downstream analyses.

### Benchmarking Metrics

We centred each normalised marker from different methods by subtracting its marker-specific mean prior to conducting Principal Component Analysis (PCA) for calculating the Silhouette coefficients and the Rozeboom vector correlation^15^ for the examination of biological and unwanted (plate) variation.

### Assessing the biological and the unwanted variation in the normalised data

#### Cluster analysis

*MAPFX* uses the PhenoGraph algorithm^8^ for its cluster analysis, and we used the same algorithm to find biological clusters for normalised data from different normalisation methods. We identified clusters based on both the normalised backbone data and the completed data matrix. Since the optimal number of clusters under different normalisation methods varied and the cluster labelling was arbitrary, we used confusion matrices to identify and match the common biological sub-populations that were found by using the normalised datasets from the three methods (we call these clusters consensus sub-populations). In this way, the cells which were grouped in the same cluster but have a different cluster number from the algorithm can be identified, and the cluster labelling can then be unified (Supp. Fig. 2).

#### Silhouette coefficients

To assess both biological and unwanted variation in the normalised data, we calculated Silhouette coefficients by using either the biological consensus labelling, or the plate annotation. Below is the formula for calculating the Silhouette coefficient:^16^

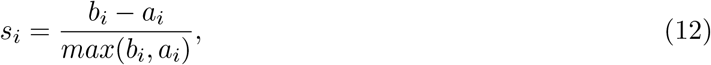

where *s*_*i*_ is the Silhouette coefficient for cell *i, a*_*i*_ is the average intra-cluster distance (i.e., average dissimilarity of cell *i* to all other cells that belong to the same cluster A), and *b*_*i*_ is the minimum of the average inter-cluster distances (i.e., the minimum of the average dissimilarity of cell *i* to all cells that belong to any other cluster C (*C* ≠ *A*). The cluster labelling can be either biological or plate annotation, depending on which aspect we are assessing. A high Silhouette coefficient indicates that the cluster labels captured the variation of the normalised data well.

#### Rozeboom correlation

We used the Rozeboom correlation^15^ (*R*_*X,Y*_) to assess the correlation between PCs (*X*), representing the normalised data, and either the biological or the plate variables (*Y*). The Rozeboom correlation is obtained by the following formula:

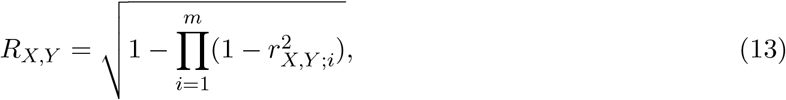

where 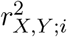 is the *i*^*th*^ nonzero canonical correlation between a PC from the set *X* and a dummy variable of either (1) biological or (2) plate variables from set *Y*, and *i* = 1, …, *m*. A higher Rozeboom correlation represents a stronger linear correlation between the normalised data and either the biological or the plate variables.

### The goodness-of-fit of the imputation models

We used the *R*^2^ statistic to quantify the distance between the measurement (the observed value) and the imputation (the predicted value).

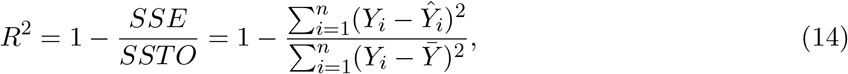

where *Y*_*i*_ is the observed value, 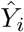 is the imputed value, and 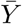 is the mean value of the observed value calculated from *n* cells.

### Assessing contribution of the protein markers to cell type refinement

We used the *F*-statistic to quantify the contribution of every protein marker to the cell type refinement and ranked their value with each other. For the cells belonging to the same broad type, we calculated the *F*-statistic for protein *p* with the following formula.

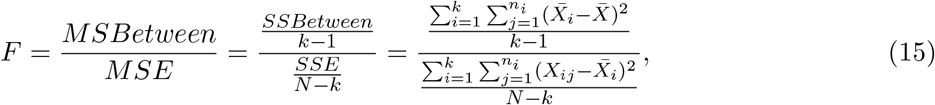

where *MSBetween* is the between-subtype variance, *MSE* is the within-subtype variance, *k* is the number of subtypes that can be found by using the completed data, *n*_*i*_ is the number of cells that are from subtype *i, N* is the number of cells that belong to a particular broad cell type, and *X*_*ij*_ is the protein intensity of cell *j* from the subtype *i*, 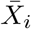 is the mean intensity of subtype *i*, 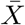 is the overall intensity calculated from the cells belonging to this broad cell type. The protein intensities are on the natural logarithm scale. The markers with a higher *F* value are the markers that contribute to the differentiation of subtypes more than the ones with a lower *F* value.

## Results

We used the publicly available dataset from Becht *et al*. (2021)^4^ to motivate our method development. The dataset contains twenty-eight million single cells that were extracted from tissues of murine lungs at the steady-state, and the majority of the cells are immune cells. After staining with 14 fluorescently labelled antibodies (14 backbone markers) those cells were allocated to three 96-well plates (266 wells with samples, 22 wells in plate 3 left blank) with most of the wells contained one unique lyophilised infinity antibody. The list of the backbone markers and the infinity markers is shown in the supplementary material (Supp. Table 1). To conduct normalisation and imputation, we used the same 2.66 million cells that were sampled by Becht *et al*., from their 28 million cells.

### The need for data normalisation

#### Evidence of background noise

Eleven isotype controls were allocated to 11 wells in the Becht *et al*. (2021) dataset, with a different isotype control in each of the 11 wells. As there is no specific antigen binding for the isotype controls, we could consider the measurements from the isotype control wells to be noise. To examine the existence of background noise, we produced histograms of the measurements from those wells. These measurements are centred around zero in all the eleven wells (Fig. 2A), which provides support for using the symmetric Normal distribution for modelling the noise component with *mapfx*.*norm*.

**Figure 2.**
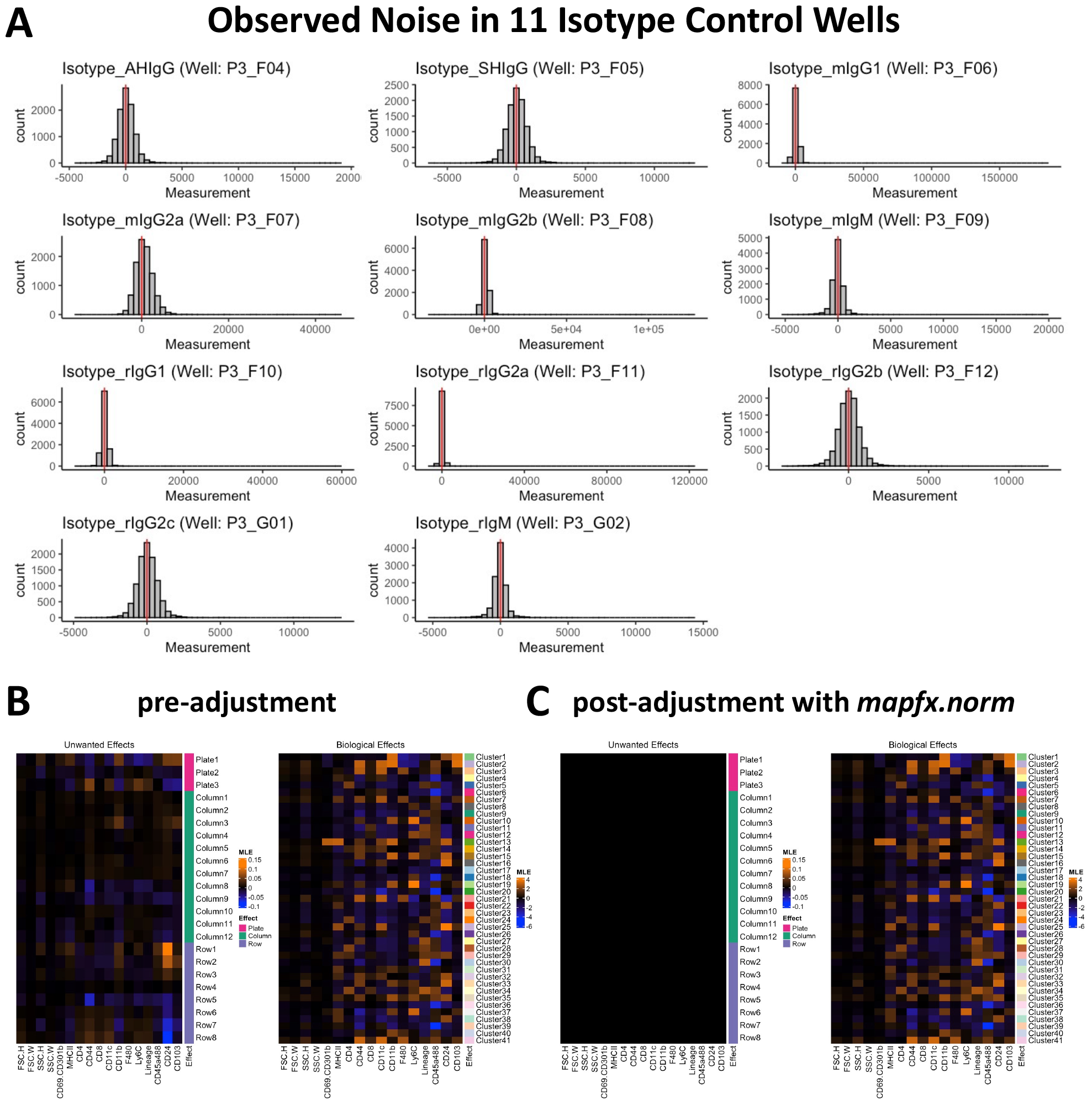
Noise, and the unwanted and the biological effects in the data. (A) Observed noise in the wells containing isotype controls (raw measurements) with a vertical line at zero in each histogram. (B) Maximum likelihood estimates of the unwanted and the biological effects estimated from the data before (pre-adjustment) and after (post-adjustment) removal of unwanted variation with *mapfx*.*norm*. Orange represents a positive effect, whereas blue indicates a negative effect. The colour-key is fixed for both the pre- and post- results. *mapfx*.*norm* managed to remove unwanted effects from the data while preserving biological effects.

#### Well effect in the background corrected data

We fitted log-linear models (equation 6) to our background-corrected data and estimated the biological and the unwanted plate, column, and row effects. Unwanted effects do indeed exist in the unadjusted data (Fig. 2B), but *mapfx*.*norm* managed to greatly mitigate the plate, column, and row effects while preserving biological effects in the data (Fig. 2C).

### Benchmarking normalisation methods

#### Biological and unwanted variation in the normalised datasets

For investigating and comparing the biology revealed by the completed datasets under different normalisation methods, we used consensus clusters for the assessment. To ease computational burdens, we performed random sampling stratified by the consensus clusters to extract 100 cells from each well, giving 26,600 cells in total for the following assessment. First, we assessed the retention of biological signal in the normalised data. The top two PCs from *mapfx*.*norm* normalised data have the strongest correlation with biological factors (0.997) among the three methods, and it remains the highest throughout (Fig. 3A). Similarly, *mapfx*.*norm* leads to the highest mean Silhouette scores across different numbers of cumulative PCs with 0.395 at PC1:10 which is the highest (Fig. 3C). Overall, *mapfx*.*norm* preserved more biological variation than alternative methods. Second, when examining the unwanted variation in the normalised data, *mapfx*.*norm* generally removed batch (plate) effect as much as the other methods, with less than 0.01 numerical difference compared to *lgc*.*z* and *lgc*.*comb*.*bio* (Fig. 3B and 3D). Moreover, *mapfx*.*norm* could outperform the alternatives when we calculated Rozeboom correlations between PCs and the batch factor based on the samples stratified by their biological clusters (Supp. Fig. 3). For example, cluster C8 at PC1:10, the correlations are 0.26, 0.27, and 0.21 for *lgc*.*z, lgc*.*comb*.*bio*, and *mapfx*.*norm*, respectively, suggesting that *mapfx*.*norm* managed to remove more plate effects within the cluster. For all the other clusters, *mapfx*.*norm* was competitive, and performed as well as or slightly better than the two alternatives. In conclusion, *mapfx*.*norm* preserves more biological variation while removing similar amount of unwanted variation from the data, compared to *lgc*.*z* and *lgc*.*comb*.*bio* methods.

**Figure 3.**
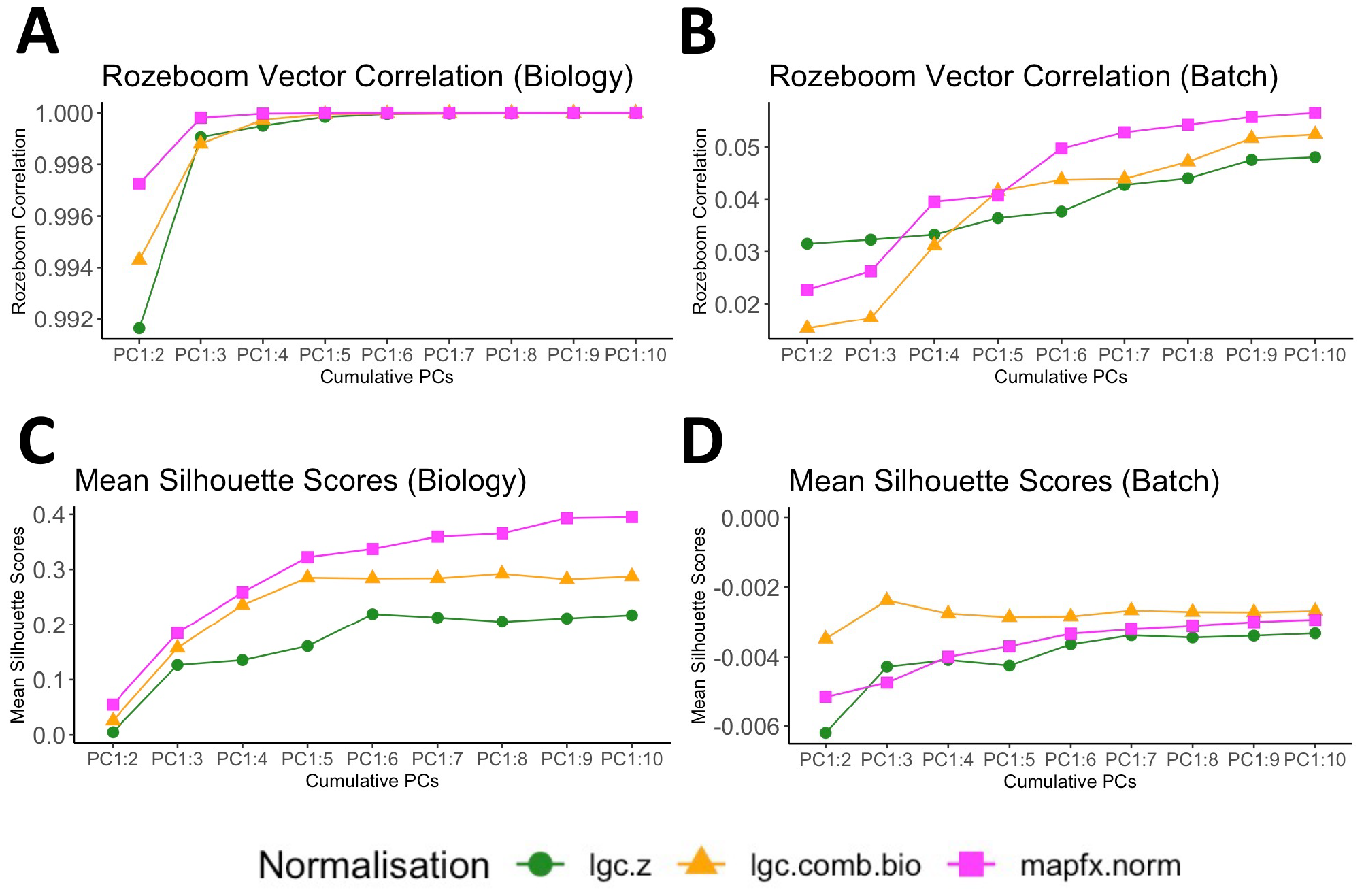
Biological and unwanted variation in the normalised data. (A) Biological and (B) batch Rozeboom correlations calculated from the three normalised data (green for *lgc*.*z*, orange for *lgc*.*comb*.*bio*, and magenta for *mapfx*.*norm*). *mapfx*.*norm* normalised data had higher correlation with biological factors started from the first two PCs while having as low correlation with batch factors (plates) as the other two methods across different numbers of cumulative PCs. Mean (C) biological and (D) batch Silhouette coefficients calculated from the three normalised data (green for *lgc*.*z*, orange for *lgc*.*comb*.*bio*, and magenta for *mapfx*.*norm*). *mapfx*.*norm* led to higher mean biological Silhouette throughout and had low mean batch Silhouette overall.

#### The goodness-of-fit of the imputation models

To examine if *mapfx*.*norm* is better than the current methods in terms of prediction of infinity markers, we used the same imputation models with the setting of the hyperparameters as addressed in Becht *et al*. (2021).^4^ The predictors were the 17 backbone markers. Analysis of 266 *R*^2^ values for the three normalisation methods showed that the median, Q3, and the maximum value of the *R*^2^ from *mapfx*.*norm* are higher than those from the alternative methods across non-linear regression models, demonstrating that the *R*^2^ values from *mapfx*.*norm* have a distribution with larger values in general, and the differences in the mean value of *R*^2^ are all statistically significant (Fig. 4A). The non-linear models (LASSO3, SVM, and XGBoost) outperformed the linear model for most of the infinity markers, indicating that the relationship between backbone markers and each infinity marker is likely to be complicated and non-linear. Additionally, among the three non-linear models, XGBoost was the fastest method, whereas LASSO3 took the longest to complete the imputation. As the overall performance is similar across different non-linear imputation models, we selected results from the XGBoost model for further investigation due to its superior computational performance. For around 80% of the infinity markers, *mapfx*.*norm* produced higher *R*^2^ values than the Logicle-based methods, indicating that its normalised data improves the accuracy of the imputations (Fig. 4B).

**Figure 4.**
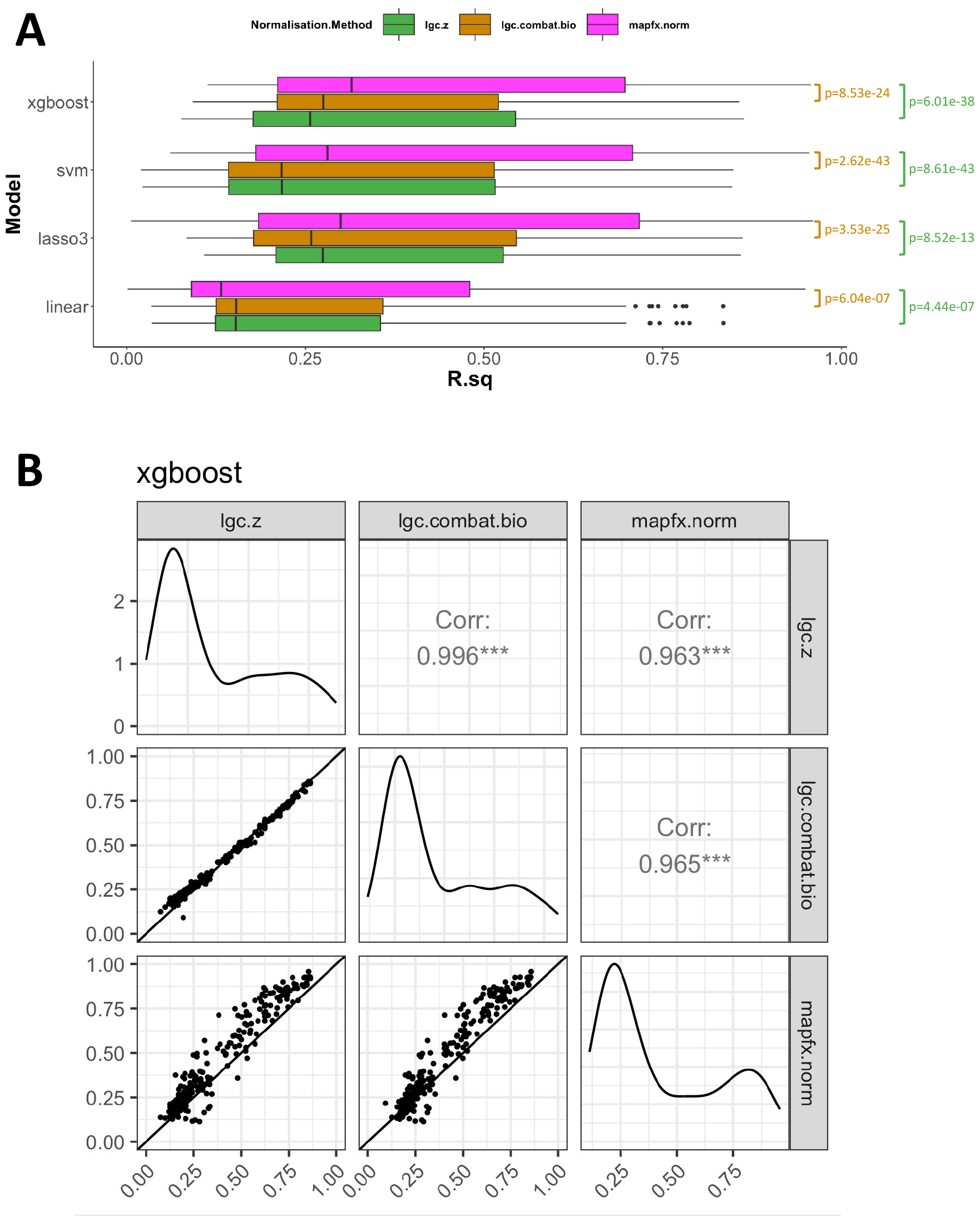
The fit of the imputation models. (A) The boxplots of 266 *R*^2^ values (horizontal axis) for three normalisation methods (colours) across four imputation models (vertical axis). *mapfx*.*norm* has higher medians *R*^2^ for non-linear models. The p-values of the paired samples T-tests (one-tailed) show that the mean values of *R*^2^ of *mapfx*.*norm* were significantly higher than the alternatives across all imputation models. (B) The pairwise scatter plots of the *R*^2^ values from the XGBoost model for the three normalisation methods. The density of *R*^2^ values is shown on the diagonal, while the lower triangular panels have the scatter plots for each pair of *R*^2^ from different methods with a 45-degree straight line, and the upper triangular panel shows the Pearson correlations of the *R*^2^ for each pair of methods. *mapfx*.*norm* provided higher *R*^2^ for around 80% of the infinity markers.

#### Refinement of cell types

General clusters (the broad cell types) were revealed by solely using informative backbone markers, and this was achieved by all normalisation methods. Applying the cell annotations from the paper by Becht *et al*. (2021),^4^ UMAP embeddings of the adjusted backbone markers were able to identify distinct populations of CD4+ T cells and B cells (Fig. 5A), however, each normalisation method produced a different degree of cell types refinement. We examined the subtypes of CD4+ T cells and B cells that were derived from the completed data matrix with imputations from the XGBoost model, using the 17 backbone markers and the 155 imputed infinity markers that did not belong to the 97 poorly imputed infinity markers claimed by Becht *et al*. (2021) (Supp. Table 2). The *R*^2^ values of these markers tend to be lower than the rest of the 155 infinity markers (Supp. Fig. 4), and were found to be unhelpful for cell type classification.

To achieve a fairer comparison for different normalisation methods, we used the broad cell types (CD4+ T and B) because they were distinctly separated, regardless of the normalisation methods (Fig. 5A). Furthermore, they had fairly large numbers of cells for examination. The numbers of cells for CD4+ T and B cells were 134,467 (10% of the whole cell population of this dataset) and 143,045 (11% of the whole cell population of this dataset), respectively. Subtypes from the completed data matrices were located distinctly in two-dimensional UMAP embeddings derived from the backbone data (Fig. 5B). The UMAP two-dimensional representation from *mapfx*.*norm* provides better refinement and separation of the subtypes. For example, the structure of the green subtype of B cells (potential immature T1 B cells) from *mapfx*.*norm* is more compact compared to Logicle-based methods. To make sure the subtypes found by *mapfx*.*norm* are meaningful but were not randomly found, we conducted MANOVA (Multivariate ANalysis Of VAriance) tests with PCs as dependent variables and the refined cell annotation from *mapfx*.*norm* as the independent variable. For B cells, the Pillai’s Trace test statistics were 1.37, 1.38, and 1.5 for *lgc*.*z, lgc*.*comb*.*bio*, and *mapfx*.*norm*, respectively. For CD4+ T cells, the corresponding Pillai’s Trace test statistics for *lgc*.*z, lgc*.*comb*.*bio*, and *mapfx*.*norm* were 1.54, 1.55, and 1.61, respectively. *mapfx*.*norm* returned higher values of the statistic for both cases suggesting that the findings are likely to be meaningful. The structure of the protein expressions for the sub-populations from *mapfx*.*norm* explicitly displays on the heatmaps formed by the top 15 key protein markers with the highest *F*-value (Fig. 5C). The markers were selected from the results of statistical analysis (F-statistic) and experts’ professional opinions.

**Figure 5.**
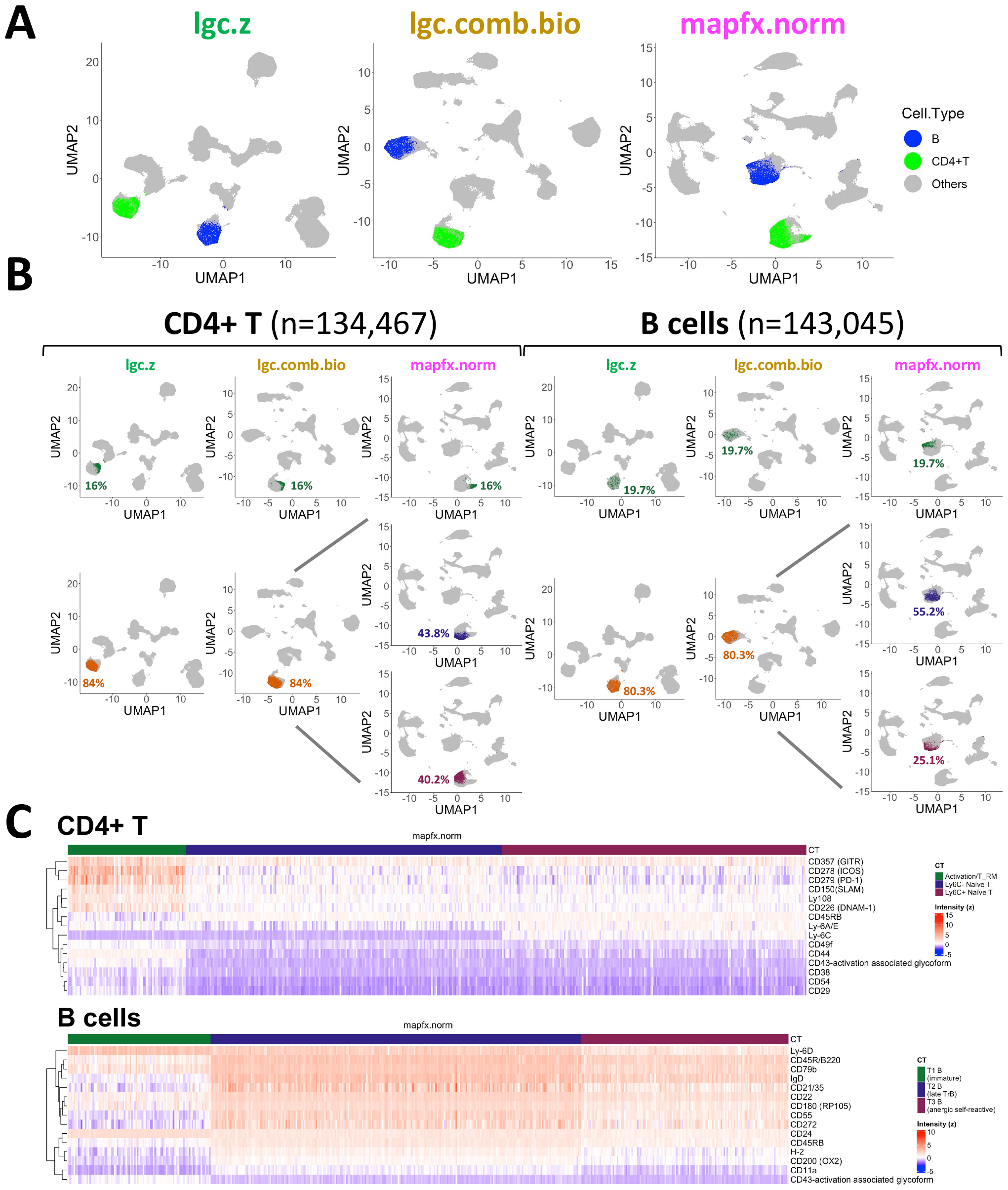
Refinement of celltypes. (A) The UMAP coordinates derived from three different normalisation methods, highlighting CD4+ T (green) and B (blue). (B) The location of the subtypes of CD4+ T and B found by each normalisation method on the corresponding UMAP coordinates and their proportion. (C) Heatmaps of a subset of cells (500 cells in each heatmap) for exhibiting the expression patterns (*mapfx*.*norm* adjusted values) of the subtypes using the key protein markers which have the highest F-values.

The subtypes found by *mapfx*.*norm* are clearly separated on the merged UMAP embeddings (Fig. 6A), indicating that the low dimensional representation reflects different protein expression levels of the subtypes. CD44 is one of the markers for differentiating Activated/Tissue-resident and Naїve CD4+ T cells, and Ly6C+ Naїve CD4+ T cells found by *mapfx*.*norm* tend to have slightly higher expression of CD44 than Ly6C-Naїve CD4+ T cells. Additionally, different stages of the B cells revealed different expression levels of CD55 marker, and the three sub-populations from *mapfx*.*norm* have different modes of the expression distribution (Fig. 6B). With the markers CD44 and CD55, the sub-populations of the CD4+ T cells and the B cells from *mapfx*.*norm* are more distinct than the representation from the alternative methods (Fig. 6C). Ly6C is the key marker that separates one sub-population of the Naїve CD4+ T cells from another (Fig. 6D). Some studies have shown that Ly6C is a marker that distinguishes high and low self-pMHC reactivity for CD4+ T cells.^17,18,19^ With other key markers from the literature^20^ (Fig. 6D), *mapfx*.*norm* has the potential to find the sub-populations of B cells - T1 (immature), T2 (late TrB), and T3 (anergic self-reactive). Moreover, in Alveolar Macrophages, another cell type that we examined, *mapfx*.*norm* found potential subtypes that could not be found by *lgc*.*z* method (Supp. Fig. 5). We found CD47 is a marker that finely addresses the difference between two sub-populations of Alveolar Macrophages. In conclusion, our proposed method *mapfx*.*norm* has the potential to provide better refinement of cell types than the Logicle-based methods. However, further experimental validation is needed to strengthen these findings.

**Figure 6.**
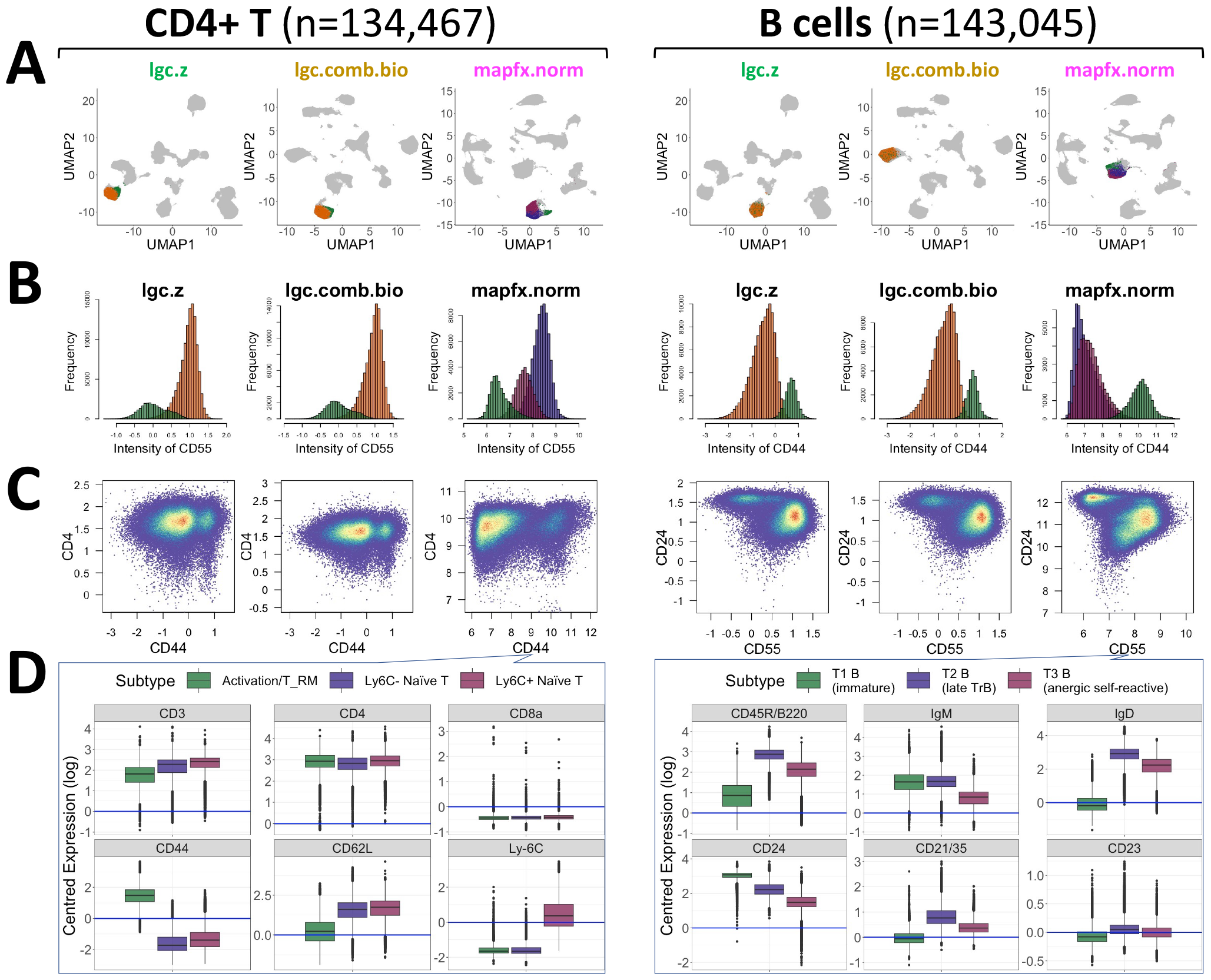
Expression level of the refined celltypes. (A) A combined version of Fig. 5B (B) The distribution of the selected imputed markers with high *F-values* for differentiating the subtypes of CD4+ T cells and B cells for each normalisation method. (C) The structure of the subtypes on the coexpression plots for each normalisation method. *mapfx*.*norm* led to clearer separation of the subtypes of both CD4+ T cells and B cells. (D) The relative expression levels (values adjusted by the protein-specific means) of the key markers for the subtypes from *mapfx*.*norm*.

#### Two additional datasets - the Intestinal data and the CD8 T Cell data

Two additional datasets were analysed to demonstrate the broad applicability of *MAPFX*. First, the Intestinal dataset^5^ contains cells from the discrete intestinal segments (duodenum, jejunum, ileum, and colon) of in-house generated Btnl2-KO and WT mice. 1,955,940 cells were allocated to 269 wells in three plates of the MPC experiment. Second, the CD8 T Cell dataset^21^ contains cells pooled from different tissues (spleen, bone marrow, liver, kidney, salivary glands, and small intestine), and the samples were enriched for CD8+ T cells. 75,986,303 cells were allocated to 269 wells in three plates of the MPC experiment. Note that we analysed the CD8 T Cell dataset based on a subset of cells (*n*=2,690,000). The lists of the backbone markers and the infinity markers for the two datasets are in the supplementary tables (Supp. Table 3 and 4). *mapfx*.*norm* removed the well effects from both datasets while retaining the biological effects (Fig. 7A). Regarding the performance of imputation, the two datasets shared the same set of infinity markers, but the Intestinal dataset had higher *R*^2^ values in general (Supp. Fig. 6) with a right-shift of its *R*^2^ distribution (Fig. 7B), suggesting that the backbone markers of the Intestinal data were more useful for the prediction of the infinity markers. Overall the Intestinal data had a higher mean value of *R*^2^ at 0.76, whereas the mean for the CD8 T Cell data is 0.57.

**Figure 7.**
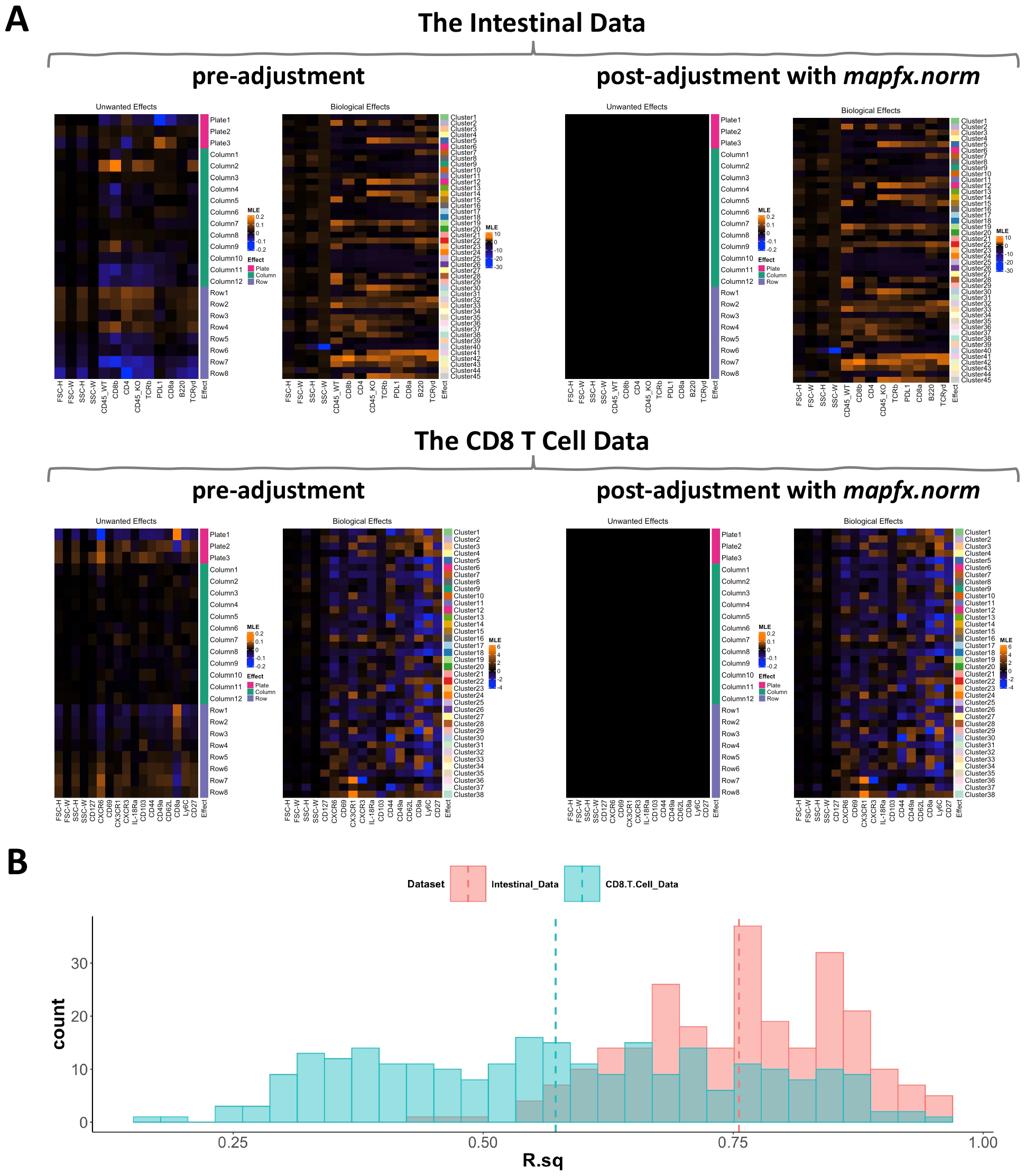
Analyses of two other datasets using our publicly available R package (*MAPFX*). (A) Maximum likelihood estimates of the unwanted and the biological effects estimated from the other two datasets before (pre-adjustment) and after (post-adjustment) removal of unwanted variation with *mapfx*.*norm*. Orange represents positive effects, whereas blue indicates negative effects. The colour-key is fixed for both pre- and post-results for each dataset. *mapfx*.*norm* managed to remove unwanted effects from the data while preserving biological effects. (B) The histogram of the 269 *R*^2^ values from the XGBoost model for the Intestinal and CD8 T Cell datasets. *R*^2^ values represent the fit of the imputation model for the infinity markers, and the infinity markers from the Intestinal dataset have higher *R*^2^ in general with a mean of 0.76, whereas the mean for the CD8 T Cell data is 0.57.

## Discussion

*MAPFX* is an end-to-end toolbox that allows us to analyse the data from MPC experiments. It starts from data pre-processing step using *mapfx*.*norm* approach followed by imputing missing infinity markers, and cluster analysis to explore the completed dataset. In addition, the way that *mapfx*.*norm* normalises the backbone markers can be directly applied to the data from the conventional fluorescence flow cytometry (FFC) assays as they essentially have the same characteristics.

Regarding the use of isotype controls for background correction, Becht *et al*. (2021) used the imputed isotype controls to remove the background noise from the data. This strategy is controversial. Assuming that there is no target for the isotype control antibodies expressed on the cells of interest can be problematic because some antibodies were chosen as isotype controls due to their unknown specificity, yet unknown specificity is not equivalent to no target. Some studies have suggested not relying on isotype controls for background correction.^22, 23, 12^ In contrast, *mapfx*.*norm* uses the small values of each protein marker to estimate the parameters of the background noise and further removes its effect from the data.

Comparing the three datasets, the Intestinal and the CD8 T Cell datasets have higher-quality imputation models, as shown by the narrower spread and the higher median of the *R*^2^ values (Supp. Fig. 8A). From the UMAP embeddings (Supp. Fig. 8B), we learnt that that the Becht *et al*. (2021) dataset has more isolated clusters. In contrast, the cluster structures of the other two datasets projected on the UMAP plots reveal fewer isolated clusters. This result suggests that the Becht *et al*. (2021) dataset has more heterogeneous and include some relatively rare cell types whose infinity markers cannot be predicted well. Conversely, the cells in the other two datasets might be more homogeneous, so it may be easier to predict the expression of the exploratory infinity markers.

Markers such as T-cell receptors (TCRs) were among the 97 infinity markers found by Becht *et al*. (2021) to be poorly imputed. It is likely that the variation of these infinity markers cannot be explained by the backbone markers. In other words, the backbone markers do not contain much information about these infinity markers. Therefore, no matter which non-linear model we used for imputation, it did not change the fact that the backbone markers are not good predictors. To improve the imputation of the infinity markers that cannot be predicted well by the backbone markers as the only predictors, we can apply the framework of Multiple Imputation by Chained Equations (MICE).^24^ In our preliminary analysis, we used the imputations from the model with backbone markers as predictors as starting values, and we updated the imputations for each infinity marker by using the backbone markers and the rest of the imputed infinity markers (Supp. Fig. 9A). This framework potentially can be used to improve the fit of the imputation models for all infinity markers using the XGBoost model (Supp. Fig. 9B). Further validation of the biology revealed by this imputation method is needed to assess the broad validity of this strategy.

Since there is always a trade-off between runtime and the accuracy of the imputations, with the *mapfx*.*norm* adjusted data, we tested the fit of the imputation models using 50%, 30%, 20%, and 10% of the cells for training the regression models. Training the models with fewer cells could save time, but doing so compromises the fit of the models (Supp. Fig. 7). Therefore, we would suggest *MAPFX* users apply at least 50% of the cells to train the imputation model to get good predictions. *MAPFX* carefully removes background noise and undesired experimental well effects with *mapfx*.*norm*, followed by imputing unmeasured infinity markers and inferring clusters using the completed dataset. In this paper, we highlighted the potential of *mapfx*.*norm* to provide better refinement of cell types than current alternatives, suggesting that our normalisation can lead to more biological insights.

## Data Availability

The *MAPFX* toolbox is implemented as a publicly available R package that is available on https://github.com/HsiaoChiLiao/MAPFX. The Becht *et al*. (2021) and the CD8 T Cell datasets are publicly available for downloads. The details are in their papers: (https://www.science.org/doi/10.1126/ sciadv.abg0505 and https://doi.org/10.1016/j.immuni.2023.06.005).^4, 21^ The Intestinal dataset is not publicly available.

## Acknowledgement

The authors would like to thank Maximilien Evrard (The Peter Doherty Institute for Infection and Immunity) and Casandra Panea (Regeneron Pharmaceuticals) for sharing their data from the LEG-ENDScreen assay. We also thank Maximilien Evrard and Phil Hodgkin (The Walter and Eliza Hall Institute) for providing their professional opinions on the biological interpretation of the cluster analyses.

## Author Contributions

A.S. and T.P.S. conceptualised the study. A.S., H.C.L, and T.P.S. designed the overall approach. H.C.L. and A.S. implemented the algorithm. H.C.L. performed data analyses and interpretation with supervision from A.S., T.P.S., and D.J.M. H.C.L. wrote the manuscript with inputs from T.P.S., A.S., and D.J.M. All authors read and approved the submitted manuscript.

## References

[1] Kalina, T., Fišer, K., Pérez-Andrés, M., Kuzílková, D., Cuenca, M., Bartol, S. J. W., Blanco, E., Engel, P., and van Zelm, M. C. (Oct, 2019) CD Maps—Dynamic Profiling of CD1–CD100 Surface Expression on Human Leukocyte and Lymphocyte Subsets. Frontiers in Immunology, 10.

[2] Labib, M. and Kelley, S. O. (Feb, 2020) Single-cell analysis targeting the proteome. Nature Reviews Chemistry, 4(3), 143–158.

[3] Papoutsoglou, G., Lagani, V., Schmidt, A., Tsirlis, K., Cabrero, D., Tegnér, J., and Tsamardinos, I. (Nov, 2019) Challenges in the Multivariate Analysis of Mass Cytometry Data: The Effect of Randomization. Cytometry Part A, 95(11), 1178–1190.

[4] Becht, E., Tolstrup, D., Dutertre, C.-A., Morawski, P. A., Campbell, D. J., Ginhoux, F., Newell, E. W., Gottardo, R., and Headley, M. B. (Sep, 2021) High-throughput single-cell quantification of hundreds of proteins using conventional flow cytometry and machine learning. Science Advances, 7(39).

[5] Panea, C., Zhang, R., VanValkenburgh, J., Ni, M., Adler, C., Wei, Y., Ochoa, F., Schmahl, J., Tang, Y., Siao, C.-J., Poueymirou, W., Espert, J., Lim, W. K., Atwal, G. S., Murphy, A. J., Sleeman, M. A., Hovhannisyan, Z., and Haxhinasto, S. (Jul, 2021) Butyrophilin-like 2 regulates site-specific adaptations of intestinal intraepithelial lymphocytes. Communications Biology, 4(1).

[6] Remšík, J., Pícková, M., Vacek, O., Fedr, R., Binó, L., Hampl, A., and Souč ek, K. (Jul, 2020) TGF-regulates Sca-1 expression and plasticity of pre-neoplastic mammary epithelial stem cells. Scientific Reports, 10(1).

[7] Weisel, N. M., Joachim, S. M., Smita, S., Callahan, D., Elsner, R. A., Conter, L. J., Chikina, M., Farber, D. L., Weisel, F. J., and Shlomchik, M. J. (Dec, 2021) Surface phenotypes of naive and memory B cells in mouse and human tissues. Nature Immunology, 23(1), 135–145.

[8] Levine, J., Simonds, E., Bendall, S., Davis, K., Amir, E.-a., Tadmor, M., Litvin, O., Fienberg, H., Jager, A., Zunder, E., Finck, R., Gedman, A., Radtke, I., Downing, J., Pe’er, D., and Nolan, G. (Jul, 2015) Data-Driven Phenotypic Dissection of AML Reveals Progenitor-like Cells that Correlate with Prognosis. Cell, 162(1), 184–197.

[9] McInnes, L., Healy, J., Saul, N., and Großberger, L. (Sep, 2018) UMAP: Uniform Manifold Approximation and Projection. Journal of Open Source Software, 3(29), 861.

[10] Parks, D. R., Roederer, M., and Moore, W. A. (2006) A new “Logicle” display method avoids deceptive effects of logarithmic scaling for low signals and compensated data. Cytometry Part A, 69A(6), 541–551.

[11] Herzenberg, L. A., Tung, J., Moore, W. A., Herzenberg, L. A., and Parks, D. R. (Jul, 2006) Interpreting flow cytometry data: a guide for the perplexed. Nature Immunology, 7(7), 681–685.

[12] Hulspas, R., O’Gorman, M. R., Wood, B. L., Gratama, J. W., and Sutherland, D. R. (Nov, 2009) Considerations for the control of background fluorescence in clinical flow cytometry. Cytometry Part B: Clinical Cytometry, 76B(6), 355–364.

[13] Johnson, W. E., Li, C., and Rabinovic, A. (Apr, 2006) Adjusting batch effects in microarray expression data using empirical Bayes methods. Biostatistics, 8(1), 118–127.

[14] Silver, J. D., Ritchie, M. E., and Smyth, G. K. (Dec, 2008) Microarray background correction: maximum likelihood estimation for the normal-exponential convolution. Biostatistics, 10(2), 352–363.

[15] Rozeboom, W. W. (Mar, 1965) Linear correlations between sets of variables. Psychometrika, 30(1), 57–71.

[16] Rousseeuw, P. J. (Nov, 1987) Silhouettes: A graphical aid to the interpretation and validation of cluster analysis. Journal of Computational and Applied Mathematics, 20, 53–65.

[17] Rogers, D., Sood, A., Wang, H., van Beek, J. J., Rademaker, T. J., Artusa, P., Schneider, C., Shen, C., Wong, D. C., Bhagrath, A., Lebel, M.-, Condotta, S. A., Richer, M. J., Martins, A. J., Tsang, J. S., Barreiro, L. B., Francois, P., Langlais, D., Melichar, H. J., and Textor, J. (Nov, 2021) Pre-existing chromatin accessibility and gene expression differences among naive CD4+ T cells influence effector potential. Cell Reports, 37(9), 110064.

[18] Guichard, V., Bonilla, N., Durand, A., Audemard-Verger, A., Guilbert, T., Martin, B., Lucas, B., and Auffray, C. (Dec, 2017) Calcium-mediated shaping of naive CD4 T-cell phenotype and function. eLife, 6, e27215.

[19] Martin, B., Auffray, C., Delpoux, A., Pommier, A., Durand, A., Charvet, C., Yakonowsky, P., de Boysson, H., Bonilla, N., Audemard, A., Sparwasser, T., Salomon, B. L., Malissen, B., and Lucas, B. (Jul, 2013) Highly self-reactive naive CD4 T cells are prone to differentiate into regulatory T cells. Nature Communications, 4(1).

[20] Zhou, Y., Zhang, Y., Han, J., Yang, M., Zhu, J., and Jin, T. (Mar, 2020) Transitional B cells involved in autoimmunity and their impact on neuroimmunological diseases. Journal of Translational Medicine, 18(1).

[21] Evrard, M., Becht, E., Fonseca, R., Obers, A., Park, S. L., Ghabdan-Zanluqui, N., Schroeder, J., Christo, S. N., Schienstock, D., Lai, J., Burn, T. N., Clatch, A., House, I. G., Beavis, P., Kallies, A., Ginhoux, F., Mueller, S. N., Gottardo, R., Newell, E. W., and Mackay, L. K. (July, 2023) Single-cell protein expression profiling resolves circulating and resident memory T cell diversity across tissues and infection contexts. Immunity, 56(7), 1664–1680.e9.

[22] Keeney, M., Gratama, J., Chin-Yee, I., and Sutherland, D. (Dec, 1998) Isotype controls in the analysis of lymphocytes and CD34+ stem and progenitor cells by flow cytometry?time to let go!. Cytometry, 34(6), 280–283.

[23] Maecker, H. T. and Trotter, J. (Sep, 2006) Flow cytometry controls, instrument setup, and the determination of positivity. Cytometry. Part A: The Journal of the International Society for Analytical Cytology, 69(9), 1037–1042.

[24] Azur, M. J., Stuart, E. A., Frangakis, C., and Leaf, P. J. (Feb, 2011) Multiple imputation by chained equations: what is it and how does it work?. International Journal of Methods in Psychiatric Research, 20(1), 40–49.

