## Supplementary Materials for "MAssively-Parallel Flow cytometry Xplorer (MAPFX): A Toolbox for Analysing Data from the Massively-Parallel Cytometry Experiments"

Hsiao-Chi Liao *et al.*

February 28, 2024

| Backbone (antigen) | Fluorochrome | Type |
|---|---|---|
| CD4 | BUV395 | preserve |
| CD44 | BUV737 | preserve |
| CD8a | BV421 | preserve |
| CD11c | BV510 | preserve |
| CD11b | BV605 | preserve |
| F480 | BV650 | preserve |
| Ly6C | BV711 | preserve |
| Lineage:<br>CD3, CD90.2, CD19,<br>CD45R/B220, Ly6G,<br>Siglec-F, NK1.1, NKp46 | BV786 | preserve |
| CD45 | Alexa Fluor 488 | preserve |
| CD103 | PerCP-Cy5-5 | preserve |
| CD24 | PE-Cy7 | preserve |
| CD69, CD301b | APC | preserve |
| MHCII | Alexa Fluor 700 | preserve |
| Live/Dead | Zombie NIR | discard |

| Backbone (physical measurement) | Description | Type |
|---|---|---|
| FSC-H | forward scatter-height | preserve |
| FSC-W | forward scatter-width | preserve |
| FSC-A | forward scatter-area | discard |
| SSC-H | side scatter-height | preserve |
| SSC-W | side scatter-width | preserve |
| SSC-A | side scatter-area | discard |

| Plate | Well | Infinity (target) | Plate | Well | Infinity (target) | Plate | Well | Infinity (target) |
|---|---|---|---|---|---|---|---|---|
| Plate1 | A01 | Blank | Plate2 | A01 | Blank.1 | Plate3 | A01 | Blank.2 |
| Plate1 | A02 | CD1 | Plate2 | A02 | CD138 | Plate3 | A02 | JAML |
| Plate1 | A03 | CD2 | Plate2 | A03 | CD140a | Plate3 | A03 | KLRG1 |
| Plate1 | A04 | CD3 | Plate2 | A04 | CD140b | Plate3 | A04 | LAP (TGF-B1) |
| Plate1 | A05 | CD4 | Plate2 | A05 | CD144 | Plate3 | A05 | LPAM-1 (Intergrin a4b7) |
| Plate1 | A06 | CD5 | Plate2 | A06 | CD146 | Plate3 | A06 | LTbR |
| Plate1 | A07 | CD6 | Plate2 | A07 | CD147 | Plate3 | A07 | Ly108 |
| Plate1 | A08 | CD8a | Plate2 | A08 | CD150(SLAM) | Plate3 | A08 | Ly-49a |
| Plate1 | A09 | CD9 | Plate2 | A09 | CD152 | Plate3 | A09 | Ly-49c/F/I/H |
| Plate1 | A10 | CD11a | Plate2 | A10 | CD153 | Plate3 | A10 | Ly49D |
| Plate1 | A11 | CD11b | Plate2 | A11 | CD154 | Plate3 | A11 | Ly49G |
| Plate1 | A12 | CD11c | Plate2 | A12 | CD155(PVR) | Plate3 | A12 | Ly49H |
| Plate1 | B01 | CD14 | Plate2 | B01 | CD157 | Plate3 | B01 | Ly-51 |
| Plate1 | B02 | CD16/32 | Plate2 | B02 | CD160 | Plate3 | B02 | Ly-6A/E |
| Plate1 | B03 | CD18 | Plate2 | B03 | CD169 | Plate3 | B03 | Ly-6C |
| Plate1 | B04 | CD19 | Plate2 | B04 | CD178 (FASL) | Plate3 | B04 | Ly-6D |
| Plate1 | B05 | CD21/35 | Plate2 | B05 | CD179 (VpreB) | Plate3 | B05 | Ly-6G |
| Plate1 | B06 | CD22 | Plate2 | B06 | CD180 (RP105) | Plate3 | B06 | GR-1 (Ly6C/Ly6G) |
| Plate1 | B07 | CD23 | Plate2 | B07 | CD183 (CXCR3) | Plate3 | B07 | Mac-2 (Galectin-3) |
| Plate1 | B08 | CD24 | Plate2 | B08 | CD193 (CCR3) | Plate3 | B08 | MAIR-IV |
| Plate1 | B09 | CD25 | Plate2 | B09 | CD194 (CCR4) | Plate3 | B09 | MD-1 |
| Plate1 | B10 | CD26 | Plate2 | B10 | CD195 (CCR5) | Plate3 | B10 | NK-1.1 |
| Plate1 | B11 | CD27 | Plate2 | B11 | CD196 (CCR6) | Plate3 | B11 | Notch1 |
| Plate1 | B12 | CD28 | Plate2 | B12 | CD197 (CCR7) | Plate3 | B12 | Notch2 |
| Plate1 | C01 | CD29 | Plate2 | C01 | CD199 (CCR9) | Plate3 | C01 | Notch3 |
| Plate1 | C02 | CD30 | Plate2 | C02 | CD200 (OX2) | Plate3 | C02 | Notch4 |
| Plate1 | C03 | CD31 | Plate2 | C03 | CD200R (OX2R) | Plate3 | C03 | PD-1H |
| Plate1 | C04 | CD34 | Plate2 | C04 | CD200R3 | Plate3 | C04 | PDC-TREM |
| Plate1 | C05 | CD36 | Plate2 | C05 | CD201 (EPCR) | Plate3 | C05 | PIR-A/B |
| Plate1 | C06 | CD38 | Plate2 | C06 | CD202b (Tie2,CD202) | Plate3 | C06 | Podoplanin |
| Plate1 | C07 | CD39 | Plate2 | C07 | CD205 | Plate3 | C07 | RAE-1g |
| Plate1 | C08 | CD40 | Plate2 | C08 | CD206(MMR) | Plate3 | C08 | RAE-1d |
| Plate1 | C09 | CD41 | Plate2 | C09 | CD207(Langerin) | Plate3 | C09 | Siglec H |
| Plate1 | C10 | CD43 | Plate2 | C10 | CD210 (IL-10R) | Plate3 | C10 | SSEA-1 |
| Plate1 | C11 | CD43-activation associated glycoform | Plate2 | C11 | CD223 (LAG3) | Plate3 | C11 | SSEA-3 |
| Plate1 | C12 | CD44 | Plate2 | C12 | CD226 (DNAM-1) | Plate3 | C12 | TCR g-d |
| Plate1 | D01 | CD45 | Plate2 | D01 | CD229 (Ly-9) | Plate3 | D01 | TCR Va11 |
| Plate1 | D02 | CD45R/B220 | Plate2 | D02 | CD252 (OX40L) | Plate3 | D02 | TCR Va2 |
| Plate1 | D03 | CD45RB | Plate2 | D03 | CD253 (TRAIL) | Plate3 | D03 | TCR Va3.2 |
| Plate1 | D04 | CD47 | Plate2 | D04 | CD254 (TRANCE, RANKL) | Plate3 | D04 | TCR Va8.3 |
| Plate1 | D05 | CD48 | Plate2 | D05 | CD255 (TWEAK) | Plate3 | D05 | TCR Vb11 |
| Plate1 | D06 | CD49a | Plate2 | D06 | CD256 (APRIL) | Plate3 | D06 | TCR Vb12 |
| Plate1 | D07 | CD49b | Plate2 | D07 | CD262 (DR5, Trail-R2) | Plate3 | D07 | TCR Vb13 |
| Plate1 | D08 | CD49b(pan-NK) | Plate2 | D08 | CD265 (RANK) | Plate3 | D08 | TCR Vb2 |
| Plate1 | D09 | CD49d | Plate2 | D09 | CD266 (FN14, TWEAK receptor) | Plate3 | D09 | TCR Vb5.1-5.2 |
| Plate1 | D10 | CD49e | Plate2 | D10 | CD267 | Plate3 | D10 | TCR Vb6 |
| Plate1 | D11 | CD49f | Plate2 | D11 | CD268 | Plate3 | D11 | TCR Vb7 |
| Plate1 | D12 | CD51 | Plate2 | D12 | CD270 (HVEM) | Plate3 | D12 | TCR Vb8.1-8.2 |
| Plate1 | E01 | CD54 | Plate2 | E01 | CD272 | Plate3 | E01 | TCR Vb9 |
| Plate1 | E02 | CD55 | Plate2 | E02 | CD273 (PD-L2) | Plate3 | E02 | TCR Vg1.1 |
| Plate1 | E03 | CD59a | Plate2 | E03 | CD274 (PD-L1) | Plate3 | E03 | TCR Vg1.1+Vg1-2 |
| Plate1 | E04 | CD61 | Plate2 | E04 | CD275 (ICOSL) | Plate3 | E04 | TCR Vg2 |
| Plate1 | E05 | CD62L | Plate2 | E05 | CD276 | Plate3 | E05 | TCR Vg3 |
| Plate1 | E06 | CD63 | Plate2 | E06 | CD278 (ICOS) | Plate3 | E06 | TCR Vd4 |
| Plate1 | E07 | CD64 | Plate2 | E07 | CD279 (PD-1) | Plate3 | E07 | TCR-b chain |
| Plate1 | E08 | CD66a | Plate2 | E08 | CD283 (TLR3) | Plate3 | E08 | TER-119 |
| Plate1 | E09 | CD68 | Plate2 | E09 | CD284 (TLR4-MD2 complex) | Plate3 | E09 | TIGIT (VSTM3( |
| Plate1 | E10 | CD69 | Plate2 | E10 | CD300f (MAIR-v) | Plate3 | E10 | Tim-1 |
| Plate1 | E11 | CD70 | Plate2 | E11 | CD309 | Plate3 | E11 | Tim-2 |
| Plate1 | E12 | CD71 | Plate2 | E12 | CD314 (NKG2D) | Plate3 | E12 | Tim-3 |
| Plate1 | F01 | CD73 | Plate2 | F01 | CD317 (BST2, PDCA1) | Plate3 | F01 | Tim-4 |
| Plate1 | F02 | CD79a | Plate2 | F02 | CD326 (EpCAM) | Plate3 | F02 | TLT-2 |
| Plate1 | F03 | CD79b | Plate2 | F03 | CD335 | Plate3 | F03 | TREM-like 4 |
| Plate1 | F04 | CD80 | Plate2 | F04 | CD351 | Plate3 | F04 | AHIgG |
| Plate1 | F05 | CD81 | Plate2 | F05 | CD355 (CRTAM) | Plate3 | F05 | SHIgG |
| Plate1 | F06 | CD83 | Plate2 | F06 | CD357 (GITR) | Plate3 | F06 | mIgG1 |
| Plate1 | F07 | CD84 | Plate2 | F07 | Allergin-1 | Plate3 | F07 | mIgG2a |
| Plate1 | F08 | CD86 | Plate2 | F08 | B7-H4 | Plate3 | F08 | mIgG2b |
| Plate1 | F09 | CD88 | Plate2 | F09 | Clec12A | Plate3 | F09 | mIgM |
| Plate1 | F10 | CD90.1 | Plate2 | F10 | Clec9A | Plate3 | F10 | rIgG1 |
| Plate1 | F11 | CD90.2 | Plate2 | F11 | CMKLR1 | Plate3 | F11 | rIgG2a |
| Plate1 | F12 | CD93 | Plate2 | F12 | CXCR7 | Plate3 | F12 | rIgG2b |
| Plate1 | G01 | CD94 | Plate2 | G01 | 33D1 | Plate3 | G01 | rIgG2c |
| Plate1 | G02 | CD96 | Plate2 | G02 | DCTRAIL-R1 | Plate3 | G02 | rIgM |
| Plate1 | G03 | CD98(4f2) | Plate2 | G03 | DCTRAIL-R2 | | | |
| Plate1 | G04 | CD103 | Plate2 | G04 | Dectin-1 | | | |
| Plate1 | G05 | CD105 | Plate2 | G05 | Delta-like1 | | | |
| Plate1 | G06 | CD106 | Plate2 | G06 | Delta-like4 | | | |
| Plate1 | G07 | CD107a (Lamp-1) | Plate2 | G07 | DR3 | | | |
| Plate1 | G08 | CD107b (Mac-3) | Plate2 | G08 | ESAM | | | |
| Plate1 | G09 | CD115 | Plate2 | G09 | F4/80 | | | |
| Plate1 | G10 | CD117 (c-kit) | Plate2 | G10 | FceR1a | | | |
| Plate1 | G11 | CD120a (TNFr Typ1-p55) | Plate2 | G11 | FR4 (Folate Receptor 4) | | | |
| Plate1 | G12 | CD120b (TNF R Type II-p75) | Plate2 | G12 | Galectin-9 | | | |
| Plate1 | H01 | CD121a (IL-1R Type I-p80) | Plate2 | H01 | GARP | | | |
| Plate1 | H02 | CD122 | Plate2 | H02 | GITR Ligand | | | |
| Plate1 | H03 | CD123 | Plate2 | H03 | H-2 | | | |
| Plate1 | H04 | CD124 | Plate2 | H04 | I-A/I-E | | | |
| Plate1 | H05 | CD126 (IL-6R a chain) | Plate2 | H05 | IFNAR-1 | | | |
| Plate1 | H06 | CD127 | Plate2 | H06 | IFNGRbeta chain | | | |
| Plate1 | H07 | CD132 | Plate2 | H07 | IgD | | | |
| Plate1 | H08 | CD133 | Plate2 | H08 | IgE | | | |
| Plate1 | H09 | CD134 (OX-40) | Plate2 | H09 | IgM | | | |
| Plate1 | H10 | CD135 | Plate2 | H10 | IL-21R | | | |
| Plate1 | H11 | CD137 | Plate2 | H11 | Integrin Beta7 | | | |
| Plate1 | H12 | CD137L (4-1BB ligand) | Plate2 | H12 | Jagged 2 | | | |

Table 1: **The backbone and the infinity panels of the dataset from Becht *et al.* (2021).**

| Plate | Well | Infinity (poorly impu.) | Plate | Well | Infinity (poorly impu.) |
|---|---|---|---|---|---|
| Plate1 | B07 | CD23 | Plate2 | F10 | Clec9A |
| Plate1 | B11 | CD27 | Plate2 | F11 | CMKLR1 |
| Plate1 | C02 | CD30 | Plate2 | F12 | CXCR7 |
| Plate1 | E11 | CD70 | Plate2 | G01 | 33D1 |
| Plate1 | F10 | CD90.1 | Plate2 | G05 | Delta-like1 |
| Plate1 | F12 | CD93 | Plate2 | G06 | Delta-like4 |
| Plate1 | G02 | CD96 | Plate2 | H02 | GITR Ligand |
| Plate1 | G09 | CD115 | Plate2 | H05 | IFNAR-1 |
| Plate1 | G11 | CD120a (TNFr Typ1-p55) | Plate2 | H06 | IFNGRbeta chain |
| Plate1 | G12 | CD120b (TNF R Type II-p75) | Plate2 | H10 | IL-21R |
| Plate1 | H01 | CD121a (IL-1R Type I-p80) | Plate2 | H12 | Jagged 2 |
| Plate1 | H03 | CD123 | Plate3 | A02 | JAML |
| Plate1 | H04 | CD124 | Plate3 | A08 | Ly-49a |
| Plate1 | H07 | CD132 | Plate3 | A11 | Ly49G |
| Plate1 | H09 | CD134 (OX-40) | Plate3 | B07 | Mac-2 (Galectin-3) |
| Plate1 | H10 | CD135 | Plate3 | B08 | MAIR-IV |
| Plate1 | H11 | CD137 | Plate3 | B11 | Notch1 |
| Plate1 | H12 | CD137L (4-1BB ligand) | Plate3 | C01 | Notch3 |
| Plate2 | A02 | CD138 | Plate3 | C03 | PD-1H |
| Plate2 | A04 | CD140b | Plate3 | C07 | RAE-1g |
| Plate2 | A07 | CD147 | Plate3 | C08 | RAE-1d |
| Plate2 | A09 | CD152 | Plate3 | C09 | Siglec H |
| Plate2 | A10 | CD153 | Plate3 | C10 | SSEA-1 |
| Plate2 | A11 | CD154 | Plate3 | C11 | SSEA-3 |
| Plate2 | B02 | CD160 | Plate3 | C12 | TCR g-d |
| Plate2 | B04 | CD178 (FASL) | Plate3 | D01 | TCR Va11 |
| Plate2 | B05 | CD179 (VpreB) | Plate3 | D02 | TCR Va2 |
| Plate2 | B09 | CD194 (CCR4) | Plate3 | D03 | TCR Va3.2 |
| Plate2 | B10 | CD195 (CCR5) | Plate3 | D04 | TCR Va8.3 |
| Plate2 | B12 | CD197 (CCR7) | Plate3 | D05 | TCR Vb11 |
| Plate2 | C05 | CD201 (EPCR) | Plate3 | D06 | TCR Vb12 |
| Plate2 | C10 | CD210 (IL-10R) | Plate3 | D07 | TCR Vb13 |
| Plate2 | C11 | CD223 (LAG3) | Plate3 | D08 | TCR Vb2 |
| Plate2 | D01 | CD229 (Ly-9) | Plate3 | D09 | TCR Vb5.1-5.2 |
| Plate2 | D02 | CD252 (OX40L) | Plate3 | D10 | TCR Vb6 |
| Plate2 | D04 | CD254 (TRANCE, RANKL) | Plate3 | D11 | TCR Vb7 |
| Plate2 | D05 | CD255 (TWEAK) | Plate3 | D12 | TCR Vb8.1-8.2 |
| Plate2 | D06 | CD256 (APRIL) | Plate3 | E01 | TCR Vb9 |
| Plate2 | D07 | CD262 (DR5, Trail-R2) | Plate3 | E02 | TCR Vg1.1 |
| Plate2 | D08 | CD265 (RANK) | Plate3 | E03 | TCR Vg1.1+Vg1-2 |
| Plate2 | D09 | CD266 (FN14, TWEAK receptor) | Plate3 | E04 | TCR Vg2 |
| Plate2 | D10 | CD267 | Plate3 | E05 | TCR Vg3 |
| Plate2 | E02 | CD273 (PD-L2) | Plate3 | E06 | TCR Vd4 |
| Plate2 | E05 | CD276 | Plate3 | E08 | TER-119 |
| Plate2 | E09 | CD284 (TLR4-MD2 complex) | Plate3 | E09 | TIGIT (VSTM3( |
| Plate2 | E11 | CD309 | Plate3 | E10 | Tim-1 |
| Plate2 | F04 | CD351 | Plate3 | E11 | Tim-2 |
| Plate2 | F07 | Allergin-1 | Plate3 | F02 | TLT-2 |
| Plate2 | F08 | B7-H4 | | | |

Table 2: **The 97 poorly imputed infinity markers claimed by Becht *et al.* (2021).**

| Backbone (antigen) | Fluorochrome | Type |
|---|---|---|
| CD45-WT | FITC-A | preserve |
| CD8b (Ly-3) | PerCP-Cy5-5-A | preserve |
| CD4 | BV510-A | preserve |
| CD45-KO | BV650-A | preserve |
| TCRb | BV711-A | preserve |
| PDL1 | APC-A | preserve |
| CD8a | Alexa Fluor 700-A | preserve |
| B220 | APC-Cy7-A | preserve |
| TCRyd (GL-3) | PE-Cy7-A | preserve |
| - | BV421-A | discard |
| - | BV605-A | discard |
| - | PE-Texas Red-A | discard |
| Live/Dead | Blue-A | discard |

| Backbone (physical measurement) | Description | Type |
|---|---|---|
| FSC-H | forward scatter-height | preserve |
| FSC-W | forward scatter-width | preserve |
| FSC-A | forward scatter-area | discard |
| SSC-H | side scatter-height | preserve |
| SSC-W | side scatter-width | preserve |
| SSC-A | side scatter-area | discard |

| Plate | Well | Infinity (target) | Plate | Well | Infinity (target) | Plate | Well | Infinity (target) |
|---|---|---|---|---|---|---|---|---|
| Plate1 | A01 | Blank | Plate2 | A01 | Blank | Plate3 | A01 | Blank |
| Plate1 | A02 | IgG Isotype Ctrl | Plate2 | A02 | CD41 | Plate3 | A02 | CD300LG |
| Plate1 | A03 | CD3ε | Plate2 | A03 | CD268 | Plate3 | A03 | CD301a |
| Plate1 | A04 | CD80 | Plate2 | A04 | CD144 | Plate3 | A04 | IL-33Rα |
| Plate1 | A05 | CD81 | Plate2 | A05 | CD370 | Plate3 | A05 | CD304 |
| Plate1 | A06 | CD154 | Plate2 | A06 | CD369 (Dectin-1, CLEC7A) | Plate3 | A06 | CD6 |
| Plate1 | A07 | Notch 1 | Plate2 | A07 | PIR-A/B | Plate3 | A07 | CD100 |
| Plate1 | A08 | CD30 | Plate2 | A08 | CD22 | Plate3 | A08 | CD104 |
| Plate1 | A09 | CD178 | Plate2 | A09 | E-Cadherin | Plate3 | A09 | CD182 |
| Plate1 | A10 | CD103 | Plate2 | A10 | CD172a (SIRPα) | Plate3 | A10 | MAdCAM-1 |
| Plate1 | A11 | Delta-like 4 | Plate2 | A11 | CD319 | Plate3 | A11 | MERTK (Mer) |
| Plate1 | A12 | CD195 | Plate2 | A12 | IgG2a, κ Isotype Ctrl | Plate3 | A12 | CD226 |
| Plate1 | B01 | Notch 4 | Plate2 | B01 | MAIR-V | Plate3 | B01 | Ly6K |
| Plate1 | B02 | CD229 (Ly-9) | Plate2 | B02 | CD146 | Plate3 | B02 | CD16/32 |
| Plate1 | B03 | CD69 | Plate2 | B03 | VISTA | Plate3 | B03 | CD150 |
| Plate1 | B04 | Notch 3 | Plate2 | B04 | CD8a | Plate3 | B04 | CD25 |
| Plate1 | B05 | JAML | Plate2 | B05 | CD275 | Plate3 | B05 | CD38 |
| Plate1 | B06 | Notch 2 | Plate2 | B06 | CD34 | Plate3 | B06 | CD133 |
| Plate1 | B07 | CD194 | Plate2 | B07 | Ly-6A/E | Plate3 | B07 | CD301b |
| Plate1 | B08 | CD152 | Plate2 | B08 | CD40 | Plate3 | B08 | CD34 |
| Plate1 | B09 | CD120a | Plate2 | B09 | CD45R/B220 | Plate3 | B09 | IgG2b, κ Isotype Ctrl |
| Plate1 | B10 | CD11c | Plate2 | B10 | CD197 | Plate3 | B10 | CD43 |
| Plate1 | B11 | Delta-like 1 | Plate2 | B11 | CD47 | Plate3 | B11 | FR4 |
| Plate1 | B12 | CD196 | Plate2 | B12 | CD98 | Plate3 | B12 | CD1d |
| Plate1 | C01 | CD29 | Plate2 | C01 | CD14 | Plate3 | C01 | CD70 |
| Plate1 | C02 | CD55 | Plate2 | C02 | CD107a (LAMP-1) | Plate3 | C02 | CD4 |
| Plate1 | C03 | Jagged 2 | Plate2 | C03 | CD18 | Plate3 | C03 | I-A/I-E |
| Plate1 | C04 | CD79b | Plate2 | C04 | Ly-6G | Plate3 | C04 | CD153 |
| Plate1 | C05 | IFN-γ R b chain | Plate2 | C05 | CD21/35 | Plate3 | C05 | CD54 |
| Plate1 | C06 | CD61 | Plate2 | C06 | Mac-2 | Plate3 | C06 | 33D1 |
| Plate1 | C07 | CD121a | Plate2 | C07 | CD199 | Plate3 | C07 | CD90.2 |
| Plate1 | C08 | TCR β chain | Plate2 | C08 | Ly-51 | Plate3 | C08 | TER-119 |
| Plate1 | C09 | FcεRIα | Plate2 | C09 | IgD | Plate3 | C09 | CD49d |
| Plate1 | C10 | CD16.2 | Plate2 | C10 | Tim-4 | Plate3 | C10 | CD24 |
| Plate1 | C11 | CD36 | Plate2 | C11 | CD71 | Plate3 | C11 | Ly-6G/Ly-6C |
| Plate1 | C12 | DcTRAIL-R1 | Plate2 | C12 | H-2 | Plate3 | C12 | CD86 |
| Plate1 | D01 | CD84 | Plate2 | D01 | CD45RB | Plate3 | D01 | CD11b |
| Plate1 | D02 | CD48 | Plate2 | D02 | CD326 | Plate3 | D02 | CD45 |
| Plate1 | D03 | CD49b | Plate2 | D03 | IgM | Plate3 | D03 | CD279 |
| Plate1 | D04 | CD120b | Plate2 | D04 | CD155 | Plate3 | D04 | RAE-1γ |
| Plate1 | D05 | CD183 | Plate2 | D05 | CD200R | Plate3 | D05 | CD8b |
| Plate1 | D06 | CD262 | Plate2 | D06 | CD254 | Plate3 | D06 | CD44 |
| Plate1 | D07 | HVEM | Plate2 | D07 | IL-21R | Plate3 | D07 | CD126 |
| Plate1 | D08 | TCR Vγ1.1 + Vγ1.2 | Plate2 | D08 | CD276 | Plate3 | D08 | CD317 |
| Plate1 | D09 | B7-H4 | Plate2 | D09 | CD9 | Plate3 | D09 | CD132 |
| Plate1 | D10 | CD339 | Plate2 | D10 | CD105 | Plate3 | D10 | CD3 |
| Plate1 | D11 | CD49a | Plate2 | D11 | CD366 | Plate3 | D11 | CD274 |
| Plate1 | D12 | PD-1H | Plate2 | D12 | 4-1BB Ligand | Plate3 | D12 | CD117 |
| Plate1 | E01 | CD85k | Plate2 | E01 | CD265 | Plate3 | E01 | CD88 |
| Plate1 | E02 | Plexin B2 | Plate2 | E02 | TLR4 (CD284)/MD2 Complex | Plate3 | E02 | CD93 |
| Plate1 | E03 | CD27 | Plate2 | E03 | CD19 | Plate3 | E03 | CD252 |
| Plate1 | E04 | DR3 | Plate2 | E04 | LPAM-1 | Plate3 | E04 | MD-1 |
| Plate1 | E05 | TCR γ/δ | Plate2 | E05 | CD62L | Plate3 | E05 | CD357 |
| Plate1 | E06 | IgG2a, κ Isotype Ctrl | Plate2 | E06 | CD23 | Plate3 | E06 | CD185 |
| Plate1 | E07 | CD45.1 | Plate2 | E07 | CD5 | Plate3 | E07 | CD37 |
| Plate1 | E08 | CD45.2 | Plate2 | E08 | CD273 | Plate3 | E08 | CD300c/d |
| Plate1 | E09 | NK-1.1 | Plate2 | E09 | CD31 | Plate3 | E09 | CD186 (CXCR6) |
| Plate1 | E10 | Ly108 | Plate2 | E10 | F4/80 | Plate3 | E10 | CD130 |
| Plate1 | E11 | CD207 | Plate2 | E11 | CD94 | Plate3 | E11 | CD198 |
| Plate1 | E12 | CX3CR1 | Plate2 | E12 | CD267 | Plate3 | E12 | CD20 |
| Plate1 | F01 | IgG1, κ Isotype Ctrl | Plate2 | F01 | Ly-49A | Plate3 | F01 | CD124 |
| Plate1 | F02 | CD66a | Plate2 | F02 | CD180 | Plate3 | F02 | IL-23R |
| Plate1 | F03 | IFNAR-1 | Plate2 | F03 | CD11a | Plate3 | F03 | CD184 |
| Plate1 | F04 | Tim-2 | Plate2 | F04 | LT beta R | Plate3 | F04 | CD2 |
| Plate1 | F05 | CD272 | Plate2 | F05 | CD122 | Plate3 | F05 | IgG2c,k Isotype Ctrl |
| Plate1 | F06 | CD64 | Plate2 | F06 | CD106 | Plate3 | F06 | Ly-6C |
| Plate1 | F07 | CD351 | Plate2 | F07 | CD365 | Plate3 | F07 | Ly-6D |
| Plate1 | F08 | LAP | Plate2 | F08 | CD115 | Plate3 | F08 | IgM, κ Isotype Ctrl |
| Plate1 | F09 | TIGIT | Plate2 | F09 | CD140a | Plate3 | F09 | CD49b |
| Plate1 | F10 | Trem-like 4 | Plate2 | F10 | PDC-TREM | Plate3 | F10 | GL7 |
| Plate1 | F11 | CD59a | Plate2 | F11 | CD135 | Plate3 | F11 | IgG Isotype Ctrl |
| Plate1 | F12 | Ly49H | Plate2 | F12 | CD127 | Plate3 | F12 | CD28 |
| Plate1 | G01 | CD90.1 | Plate2 | G01 | CD140b | Plate3 | G01 | Podoplanin |
| Plate1 | G02 | IgG2b, κ Isotype Ctrl | Plate2 | G02 | ESAM | Plate3 | G02 | CD137 |
| Plate1 | G03 | CD157 | Plate2 | G03 | CD200 | Plate3 | G03 | CD278 |
| Plate1 | G04 | CD159a | Plate2 | G04 | CD309 | Plate3 | G04 | KLRG1 |
| Plate1 | G05 | XCR1 | Plate2 | G05 | TLT-2 | Plate3 | G05 | Ly-49C/F/I/H |
| Plate1 | G06 | IgM, κ Isotype Ctrl | Plate2 | G06 | CD253 | | | |
| Plate1 | G07 | SSEA-1 | Plate2 | G07 | CD335 | | | |
| Plate1 | G08 | IgG1, κ Isotype Ctrl | Plate2 | G08 | CD205 | | | |
| Plate1 | G09 | Ig light chain κ | Plate2 | G09 | Galectin-9 | | | |
| Plate1 | G10 | Siglec H | Plate2 | G10 | CD200R3 | | | |
| Plate1 | G11 | CD255 | Plate2 | G11 | MAIR-IV | | | |
| Plate1 | G12 | CD202b | Plate2 | G12 | Ly49D | | | |
| Plate1 | H01 | GITR Ligand | Plate2 | H01 | CD123 | | | |
| Plate1 | H02 | CD147 | Plate2 | H02 | CD355 | | | |
| Plate1 | H03 | CD73 | Plate2 | H03 | CD169 | | | |
| Plate1 | H04 | CD51 | Plate2 | H04 | CD138 | | | |
| Plate1 | H05 | NKG2D | Plate2 | H05 | CD160 | | | |
| Plate1 | H06 | CD96 | Plate2 | H06 | CD39 | | | |
| Plate1 | H07 | Integrin β7 | Plate2 | H07 | GARP | | | |
| Plate1 | H08 | CD210 | Plate2 | H08 | CD179a | | | |
| Plate1 | H09 | CD83 | Plate2 | H09 | CD371 | | | |
| Plate1 | H10 | Mac-3 | Plate2 | H10 | CD63 | | | |
| Plate1 | H11 | CD223 | Plate2 | H11 | CD49e | | | |
| Plate1 | H12 | CD134 | Plate2 | H12 | CD193 | | | |

Table 3: **The backbone and the infinity panels of the Intestinal Data.**

| Backbone (antigen) | Fluorochrome | Type |
|---|---|---|
| CD127 | FJComp-450_40 Violet-A | preserve |
| CXCR6 | FJComp-530_30 Blue-A | preserve |
| CD69 | FJComp-605_40 Violet-A | preserve |
| CX3CR1 | FJComp-610_20 Yellow-A | preserve |
| CXCR3 | FJComp-660_20 Violet-A | preserve |
| IL-18Ra | FJComp-670_14 Red-A | preserve |
| CD103 | FJComp-695_40 Blue-A | preserve |
| CD49a | FJComp-710_50 Violet-A | preserve |
| CD62L | FJComp-730_45 Red-A | preserve |
| CD8a | FJComp-740_35 UV-A | preserve |
| Ly6C | FJComp-780_60 Violet-A | preserve |
| CD27 | FJComp-780_60 Yellow-A | preserve |
| CD44 | FJComp-525_50 Violet-A | Δ (not used for prediction as it's contaminated) |
| barcode | FJComp-515_30 UV-A | discard |
| barcode | FJComp-379_28 UV-A | discard |
| barcode | FJComp-820_60 UV-A (Zombie NIR) | discard |
| Live/Dead | FJComp-780_60 Red-A | discard |

| Backbone (physical measurement) | Description | Type |
|---|---|---|
| FSC-H | forward scatter-height | preserve |
| FSC-W | forward scatter-width | preserve |
| FSC-A | forward scatter-area | discard |
| SSC-H | side scatter-height | preserve |
| SSC-W | side scatter-width | preserve |
| SSC-A | side scatter-area | discard |

| Plate | Well | Infinity (target) | Plate | Well | Infinity (target) | Plate | Well | Infinity (target) |
|---|---|---|---|---|---|---|---|---|
| Plate1 | A01 | Blank | Plate2 | A01 | Blank | Plate3 | A01 | Blank |
| Plate1 | A02 | Arm_H_IgG_isotype | Plate2 | A02 | CD41 | Plate3 | A02 | CD300LG |
| Plate1 | A03 | CD3e | Plate2 | A03 | CD268 | Plate3 | A03 | CD301 |
| Plate1 | A04 | CD80 | Plate2 | A04 | CD144 | Plate3 | A04 | IL-33Ra |
| Plate1 | A05 | CD81 | Plate2 | A05 | CD370 | Plate3 | A05 | CD304 |
| Plate1 | A06 | CD154 | Plate2 | A06 | CD369 | Plate3 | A06 | CD6 |
| Plate1 | A07 | Notch1 | Plate2 | A07 | PIR-A_B | Plate3 | A07 | CD100 |
| Plate1 | A08 | CD30 | Plate2 | A08 | CD22 | Plate3 | A08 | CD104 |
| Plate1 | A09 | CD178 | Plate2 | A09 | CD324 | Plate3 | A09 | CXCR2 |
| Plate1 | A10 | CD103 | Plate2 | A10 | CD172a | Plate3 | A10 | MadCAM-1 |
| Plate1 | A11 | DLL4 | Plate2 | A11 | CD319 | Plate3 | A11 | MERTK |
| Plate1 | A12 | CCR5 | Plate2 | A12 | Rat_IgG2a_isotype | Plate3 | A12 | CD226 |
| Plate1 | B01 | Notch4 | Plate2 | B01 | MAIR-V | Plate3 | B01 | Ly6K |
| Plate1 | B02 | CD229 | Plate2 | B02 | CD146 | Plate3 | B02 | CD16-32 |
| Plate1 | B03 | CD69 | Plate2 | B03 | VISTA | Plate3 | B03 | CD150 |
| Plate1 | B04 | Notch3 | Plate2 | B04 | CD8a | Plate3 | B04 | CD25 |
| Plate1 | B05 | JAML | Plate2 | B05 | CD275 | Plate3 | B05 | CD38 |
| Plate1 | B06 | Notch2 | Plate2 | B06 | CD34 | Plate3 | B06 | CD133 |
| Plate1 | B07 | CCR4 | Plate2 | B07 | Ly6A-E | Plate3 | B07 | CD301b |
| Plate1 | B08 | CD152 | Plate2 | B08 | CD40 | Plate3 | B08 | CD34 |
| Plate1 | B09 | CD120a | Plate2 | B09 | B220 | Plate3 | B09 | Rat_IgG2b_isotype |
| Plate1 | B10 | CD11c | Plate2 | B10 | CCR7 | Plate3 | B10 | CD43 |
| Plate1 | B11 | DLL1 | Plate2 | B11 | CD47 | Plate3 | B11 | FR4 |
| Plate1 | B12 | CCR6 | Plate2 | B12 | CD98 | Plate3 | B12 | CD1d |
| Plate1 | C01 | CD29 | Plate2 | C01 | CD14 | Plate3 | C01 | CD70 |
| Plate1 | C02 | CD55 | Plate2 | C02 | CD107a | Plate3 | C02 | CD4 |
| Plate1 | C03 | Jagged2 | Plate2 | C03 | CD18 | Plate3 | C03 | MHC-II_IA-IE |
| Plate1 | C04 | CD79b | Plate2 | C04 | Ly6G | Plate3 | C04 | CD153 |
| Plate1 | C05 | IFNgRb | Plate2 | C05 | CD21-35 | Plate3 | C05 | CD54 |
| Plate1 | C06 | CD61 | Plate2 | C06 | Mac-2 | Plate3 | C06 | 33D1 |
| Plate1 | C07 | CD121a | Plate2 | C07 | CCR9 | Plate3 | C07 | CD90-2 |
| Plate1 | C08 | TCRb | Plate2 | C08 | Ly-51 | Plate3 | C08 | Ter119 |
| Plate1 | C09 | FceR1a | Plate2 | C09 | IgD | Plate3 | C09 | CD49d |
| Plate1 | C10 | CD16-2 | Plate2 | C10 | Tim-4 | Plate3 | C10 | CD24 |
| Plate1 | C11 | CD36 | Plate2 | C11 | CD71 | Plate3 | C11 | Gr1 |
| Plate1 | C12 | DcTRAILR1 | Plate2 | C12 | H-2 | Plate3 | C12 | CD86 |
| Plate1 | D01 | CD84 | Plate2 | D01 | CD45RB | Plate3 | D01 | CD11b |
| Plate1 | D02 | CD48 | Plate2 | D02 | CD326 | Plate3 | D02 | CD45 |
| Plate1 | D03 | CD49b | Plate2 | D03 | IgM | Plate3 | D03 | CD279 |
| Plate1 | D04 | CD120b | Plate2 | D04 | CD155 | Plate3 | D04 | RAE-1g |
| Plate1 | D05 | CXCR3 | Plate2 | D05 | CD200R | Plate3 | D05 | CD8b |
| Plate1 | D06 | CD262 | Plate2 | D06 | CD254 | Plate3 | D06 | CD44 |
| Plate1 | D07 | HVEM | Plate2 | D07 | IL-21R | Plate3 | D07 | CD126 |
| Plate1 | D08 | TCRVg11-12 | Plate2 | D08 | CD276 | Plate3 | D08 | CD317 |
| Plate1 | D09 | B7-H4 | Plate2 | D09 | CD9 | Plate3 | D09 | CD132 |
| Plate1 | D10 | CD339 | Plate2 | D10 | CD105 | Plate3 | D10 | CD3 |
| Plate1 | D11 | CD49a | Plate2 | D11 | CD366 | Plate3 | D11 | CD274 |
| Plate1 | D12 | PD-1H | Plate2 | D12 | 4-1BB-ligand | Plate3 | D12 | CD117 |
| Plate1 | E01 | CD85k | Plate2 | E01 | CD265 | Plate3 | E01 | CD88 |
| Plate1 | E02 | PlexinB2 | Plate2 | E02 | TLR4-MD2 | Plate3 | E02 | CD93 |
| Plate1 | E03 | CD27 | Plate2 | E03 | CD19 | Plate3 | E03 | CD252 |
| Plate1 | E04 | DR3 | Plate2 | E04 | a4b7_integrin | Plate3 | E04 | MD-1 |
| Plate1 | E05 | TCRgd | Plate2 | E05 | CD62L | Plate3 | E05 | CD357 |
| Plate1 | E06 | Mouse_IgG2a_isotype | Plate2 | E06 | CD23 | Plate3 | E06 | CXCR5 |
| Plate1 | E07 | CD45-1 | Plate2 | E07 | CD5 | Plate3 | E07 | CD37 |
| Plate1 | E08 | CD45-2 | Plate2 | E08 | CD273 | Plate3 | E08 | CD300c-d |
| Plate1 | E09 | NK11 | Plate2 | E09 | CD31 | Plate3 | E09 | CXCR6 |
| Plate1 | E10 | Ly108 | Plate2 | E10 | F4-80 | Plate3 | E10 | CD130 |
| Plate1 | E11 | CD207 | Plate2 | E11 | CD94 | Plate3 | E11 | CCR8 |
| Plate1 | E12 | CX3CR1 | Plate2 | E12 | CD267 | Plate3 | E12 | CD20 |
| Plate1 | F01 | Mouse_IgG1_isotype | Plate2 | F01 | Ly-49A | Plate3 | F01 | CD124 |
| Plate1 | F02 | CD66a | Plate2 | F02 | CD180 | Plate3 | F02 | IL-23R |
| Plate1 | F03 | IFNAR-1 | Plate2 | F03 | CD11a | Plate3 | F03 | CXCR4 |
| Plate1 | F04 | Tim-2 | Plate2 | F04 | LTbR | Plate3 | F04 | CD2 |
| Plate1 | F05 | CD272 | Plate2 | F05 | CD122 | Plate3 | F05 | Rat_IgG2c_isotype |
| Plate1 | F06 | CD64 | Plate2 | F06 | CD106 | Plate3 | F06 | Ly6C |
| Plate1 | F07 | CD351 | Plate2 | F07 | CD365 | Plate3 | F07 | Ly6D |
| Plate1 | F08 | LAP | Plate2 | F08 | CD115 | Plate3 | F08 | Rat_IgM_isotype |
| Plate1 | F09 | TIGIT | Plate2 | F09 | CD140a | Plate3 | F09 | CD49b |
| Plate1 | F10 | Trem-like4 | Plate2 | F10 | PDC-TREM | Plate3 | F10 | GL7 |
| Plate1 | F11 | CD59a | Plate2 | F11 | CD135 | Plate3 | F11 | Syr_Ham_IgG_isotype |
| Plate1 | F12 | Ly49H | Plate2 | F12 | CD127 | Plate3 | F12 | CD28 |
| Plate1 | G01 | CD90-1 | Plate2 | G01 | CD140b | Plate3 | G01 | Podoplanin |
| Plate1 | G02 | Mouse_IgG2b_isotype | Plate2 | G02 | ESAM | Plate3 | G02 | CD137 |
| Plate1 | G03 | CD157 | Plate2 | G03 | CD200 | Plate3 | G03 | CD278 |
| Plate1 | G04 | CD159a | Plate2 | G04 | CD309 | Plate3 | G04 | KLRG1 |
| Plate1 | G05 | XCR1 | Plate2 | G05 | TLT-2 | Plate3 | G05 | Ly-49C-F-I-H |
| Plate1 | G06 | Mouse_IgM_isotype | Plate2 | G06 | CD253 | | | |
| Plate1 | G07 | SSEA-1 | Plate2 | G07 | CD335 | | | |
| Plate1 | G08 | Rat_IgG1_isotype | Plate2 | G08 | CD205 | | | |
| Plate1 | G09 | Ig_light_chain_k | Plate2 | G09 | Galectin-9 | | | |
| Plate1 | G10 | Siglec-H | Plate2 | G10 | CD200R3 | | | |
| Plate1 | G11 | CD255 | Plate2 | G11 | MAIR-IV | | | |
| Plate1 | G12 | CD202b | Plate2 | G12 | Ly49D | | | |
| Plate1 | H01 | GITR_Ligand | Plate2 | H01 | CD123 | | | |
| Plate1 | H02 | CD147 | Plate2 | H02 | CD355 | | | |
| Plate1 | H03 | CD73 | Plate2 | H03 | CD169 | | | |
| Plate1 | H04 | CD51 | Plate2 | H04 | CD138 | | | |
| Plate1 | H05 | NKG2D | Plate2 | H05 | CD160 | | | |
| Plate1 | H06 | CD96 | Plate2 | H06 | CD39 | | | |
| Plate1 | H07 | IntegrinB7 | Plate2 | H07 | GARP | | | |
| Plate1 | H08 | CD210 | Plate2 | H08 | CD179a | | | |
| Plate1 | H09 | CD83 | Plate2 | H09 | CD371 | | | |
| Plate1 | H10 | Mac-3 | Plate2 | H10 | CD63 | | | |
| Plate1 | H11 | CD223 | Plate2 | H11 | CD49e | | | |
| Plate1 | H12 | CD134 | Plate2 | H12 | CCR3 | | | |

Table 4: **The backbone and the infinity panels of the CD8 T Cell Data.**

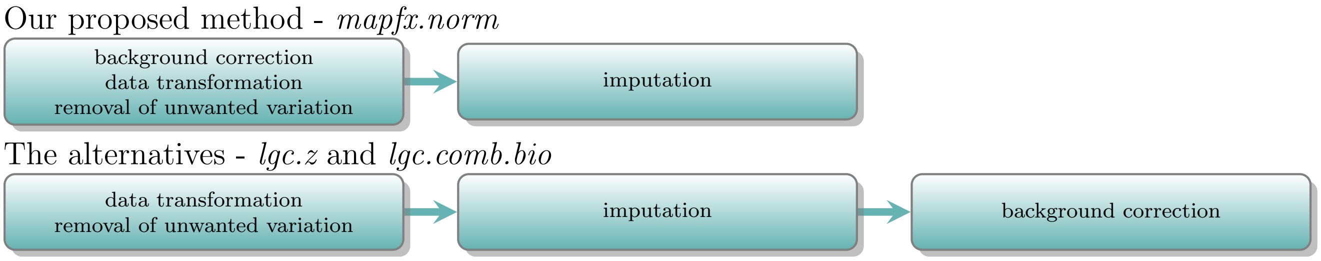


Figure 1: **The workflows of the three normalisation methods.** *mapfx.norm* performs the background correction before imputation, however, *lgc.z* and *lgc.comb.bio* correct the background noise using the imputed data.

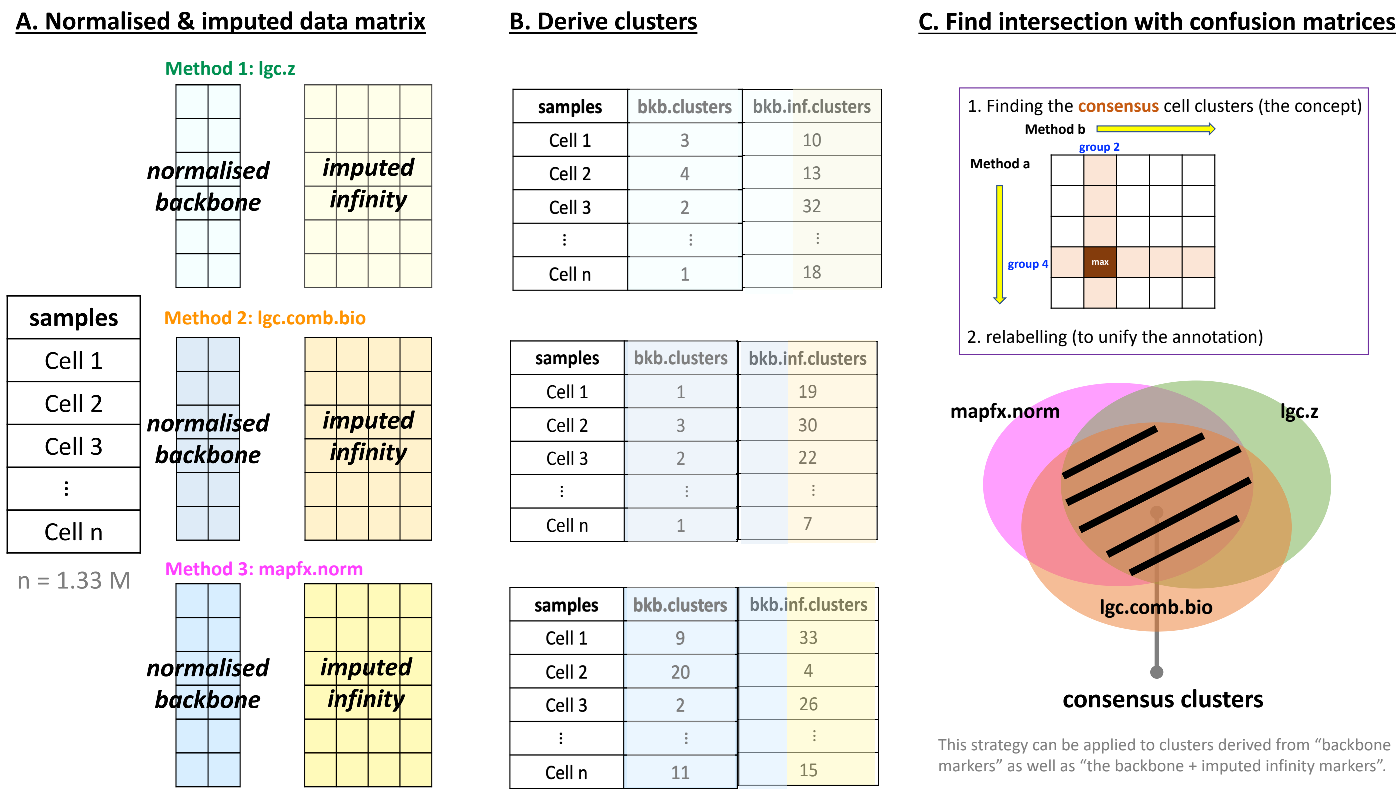


Figure 2: **The schematic diagram for the procedure of finding consensus clusters.** (A) The completed (backbone + imputed infinity) datasets from the three normalisation methods. (B) The clusters (arbitrary numbers given) derived from each completed dataset by using either normalised backbone markers (the middle column) or the completed data matrix (the right column). (C) Using the confusion matrix, the consensus clusters are the clusters that reach the row-wise and the column-wise maximum simultaneously (note that different datasets may have a different labelling for the same cluster). After finding the intersection (consensus clusters), relabelling is done to unify the cluster labels across methods.

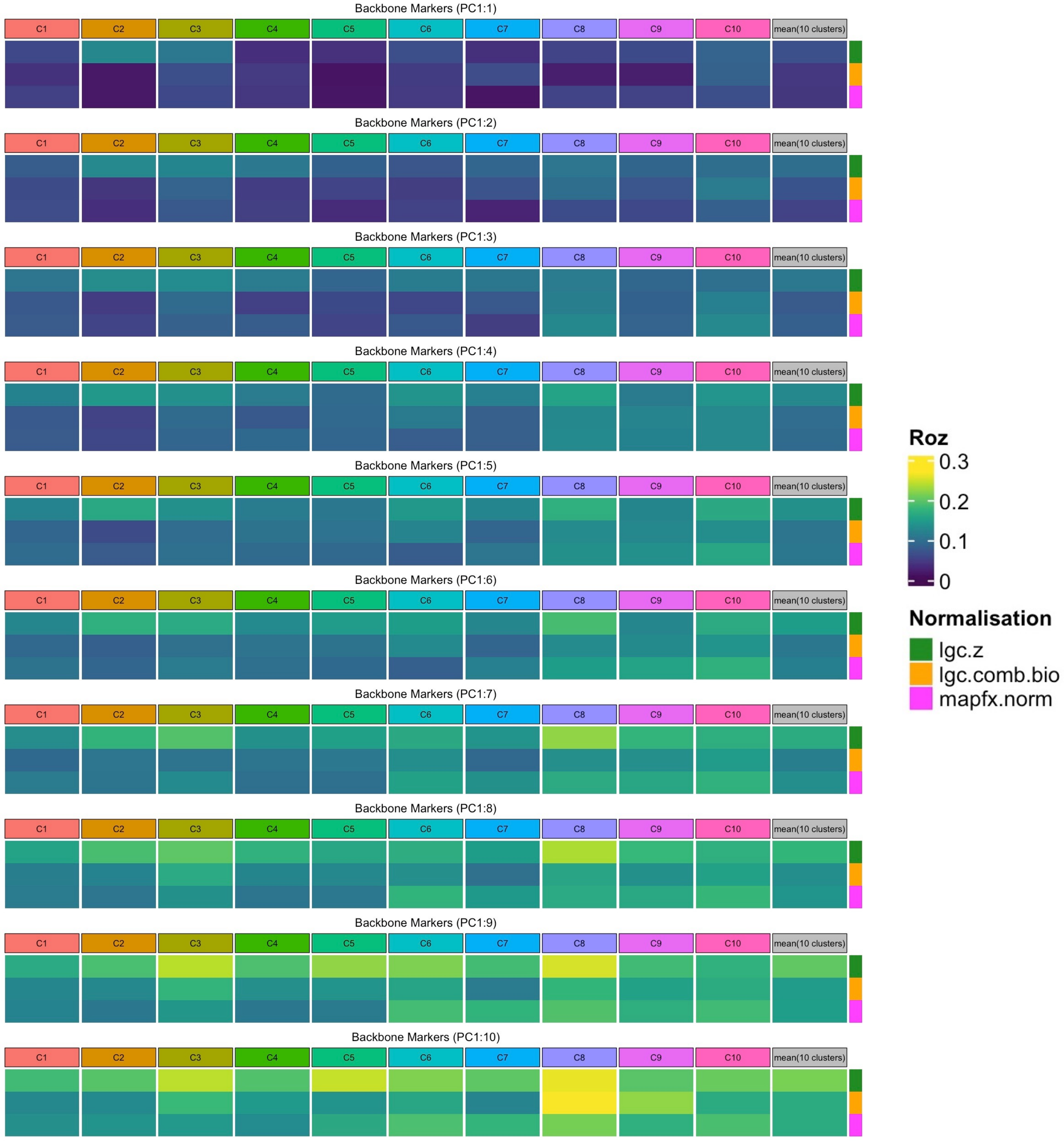


Figure 3: **The heatmaps of the Rozeboom correlations between PCs and the plate factors.** Each heatmap represents a given number of cumulative PCs used to calculate the cluster-specific canonical correlation between data (PCs) and the plate factors. In each plot, columns represent certain biological clusters (C1 is the largest cluster and C10 is the smallest cluster) with the last column (coloured in gray) indicating the mean value calculated from the 10 clusters. Rows are different normalisation methods (green for *lgc.z*, orange for *lgc.comb.bio*, and magenta for *mapfx.norm*).

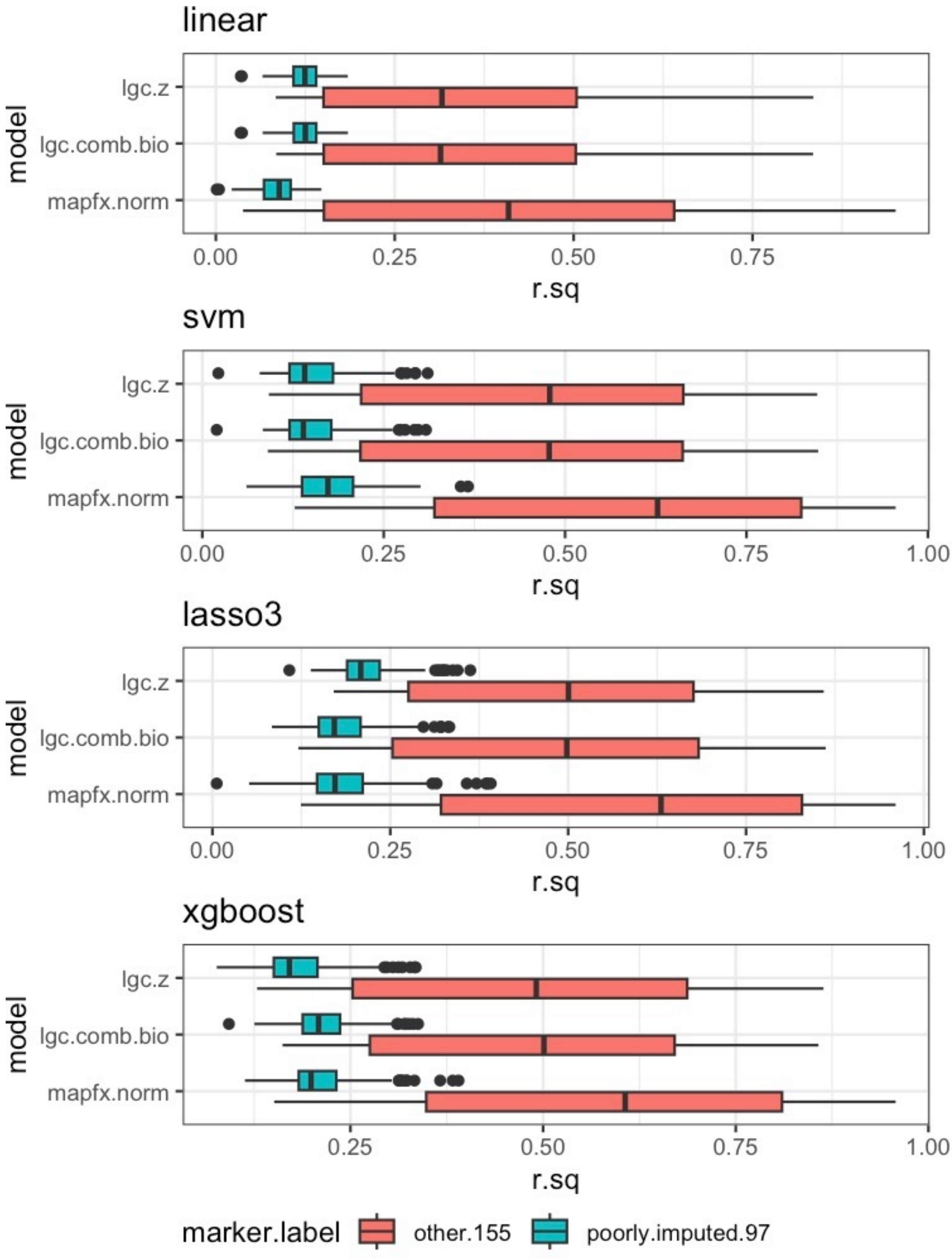


Figure 4: **The distributions of the $R^2$ of the poorly imputed (found by Becht *et al.* (2021)) and the rest of the 155 infinity markers.** The boxplots of the $R^2$ of the 97 poorly imputed infinity markers (cyan) and the rest of the 155 infinity markers (coral) from the four regression models using normalised datasets from the three methods.

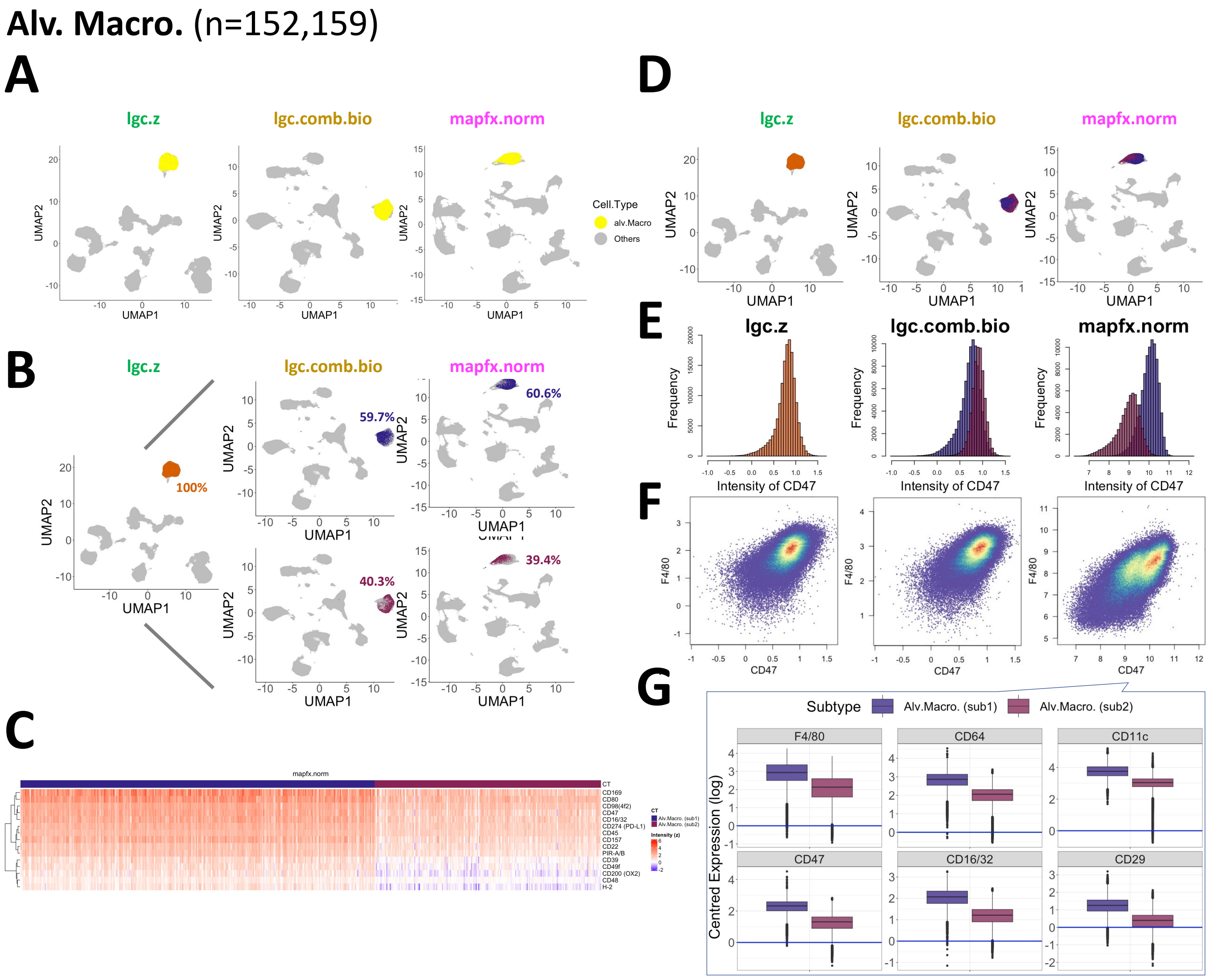


Figure 5: **Refinement of Alveolar Macrophages.** (A) The UMAP coordinates derived from three different normalisation methods, highlighting Alveolar Macrophages (yellow). (B) The location of the subtypes of Alveolar Macrophages found by each normalisation method on the corresponding UMAP coordinates and their proportion. (C) Heatmaps of a subset of cells (500 cells in the heatmap) for exhibiting the expression patterns (*mapfx.norm* adjusted values) of the subtypes using the key protein markers with the highest F-values. (D) A combined version of Supp. Fig. B (E) The distribution of the selected imputed markers for the subtypes of Alveolar Macrophages for each normalisation method. (F) The structure of the subtypes on the coexpression plots for each normalisation method. (G) The relative expression levels (values adjusted by the protein-specific means) of the key markers for the subtypes from *mapfx.norm*. CD47 is a marker that can be used to differentiate the two sub-populations.

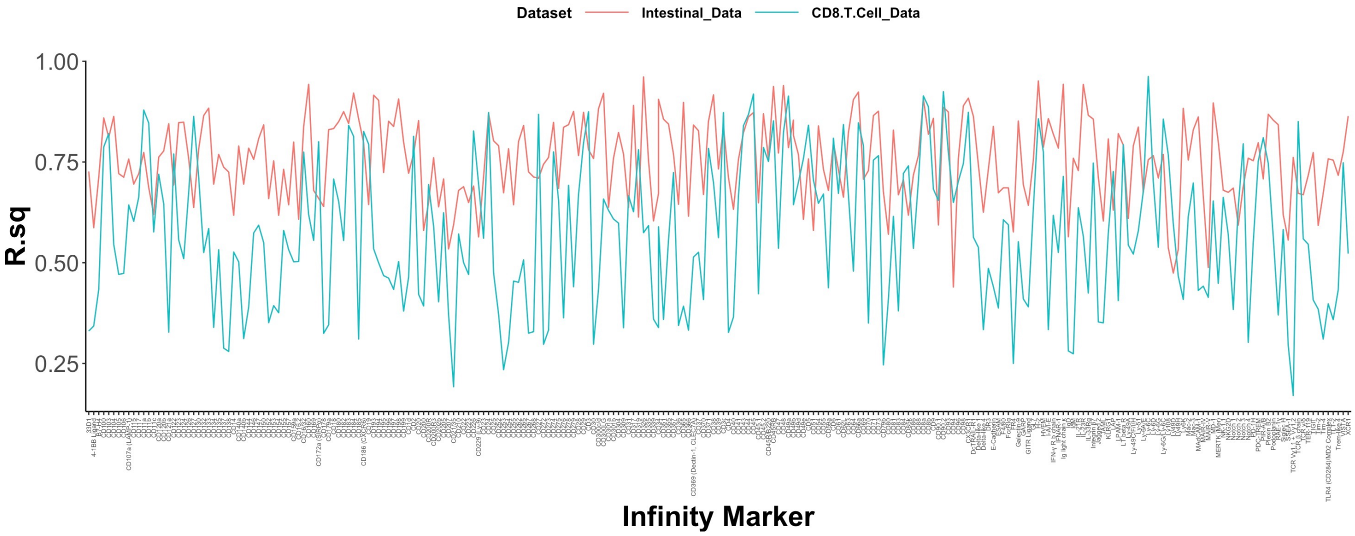


Figure 6: **A direct $R^2$ comparison for the same infinity markers (on the x-axis) between Intestinal and CD8 T Cell datasets.** The $R^2$ values from the CD8 T Cell data are mostly lower than the ones from the Intestinal data.

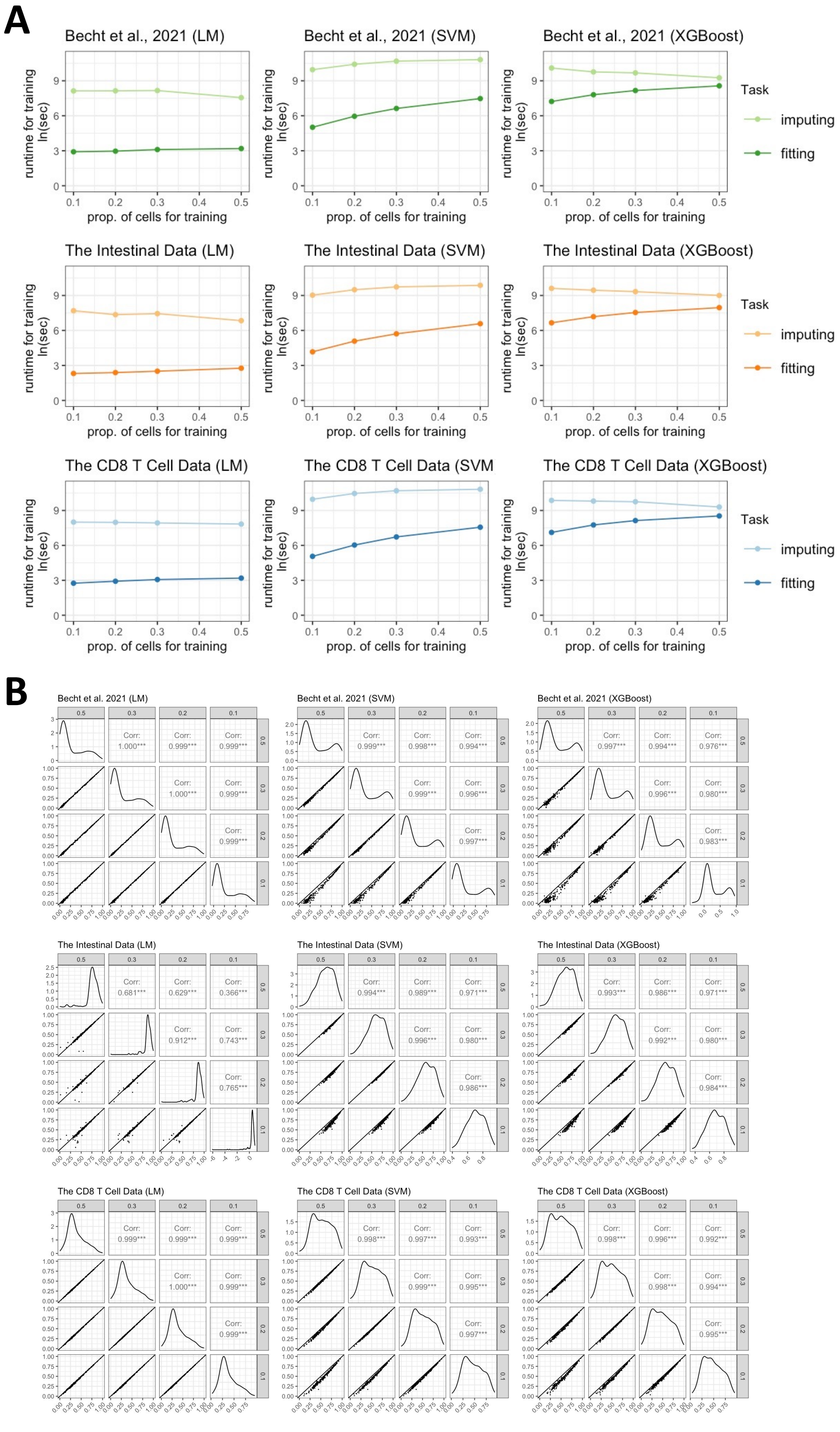


Figure 7: **The runtime and the goodness-of-fit of the imputation with different numbers of cells for training.** (A) The runtime ($log_e$(sec)) for running three regression models by using different proportion of cells (10%, 20%, 30%, and 50%) for training using three datasets (Becht et al. in green, the Intestinal Data in orange, and the CD8 T Cell Data in blue). The lines in darker colours indicate the runtime of fitting (training) the model, and the lines in lighter colours represent the runtime of imputation. (B) The pairwise plots of $R^2$ values from each regression model for the three datasets with different proportion of cells for model training. The density of $R^2$ values is shown on the diagonal, the lower triangular panel has the scatter plots for each pair of settings of the training model with a 45-degree straight line as X=Y, and the upper triangular panel shows the Pearson correlations of the $R^2$ values from each pair. More cells for training led to higher $R^2$ values.

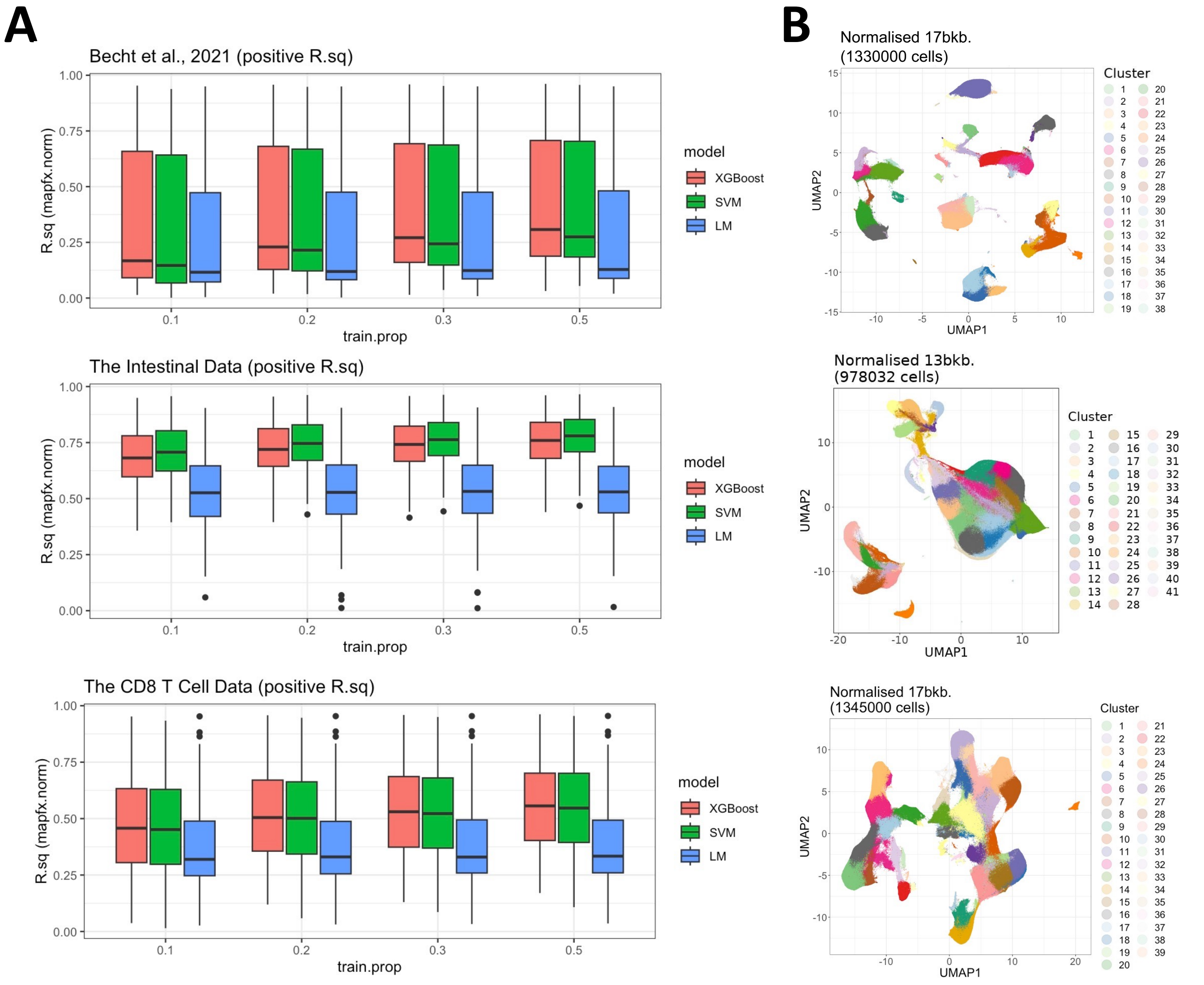


Figure 8: **The boxplots of the $R^2$ values of the infinity markers from three regression models using three datasets with different sizes of cells for model training, and the UMAP plots derived from the *mapfx.norm* adjusted backbone markers.** (A) For each panel, the y-axis indicates the magnitude of the $R^2$ values, and the x-axis is the proportion of cells used for model training. Each box contains 266 values for the Becht *et al.* (2021) dataset, and 269 values for the other two datasets. The first panel is the result from the Becht *et al.* (2021) data, while the second and the third panels are from the Intestinal and the CD8 T Cell datasets, respectively. The colours represent different regression models, coral for XGBoost, green for SVM, and blue for LM. (B) For each dataset, the UMAP coordinates were derived from the corresponding *mapfx.norm* adjusted backbone markers and coloured by the clusters found by the PhenoGraph algorithm.

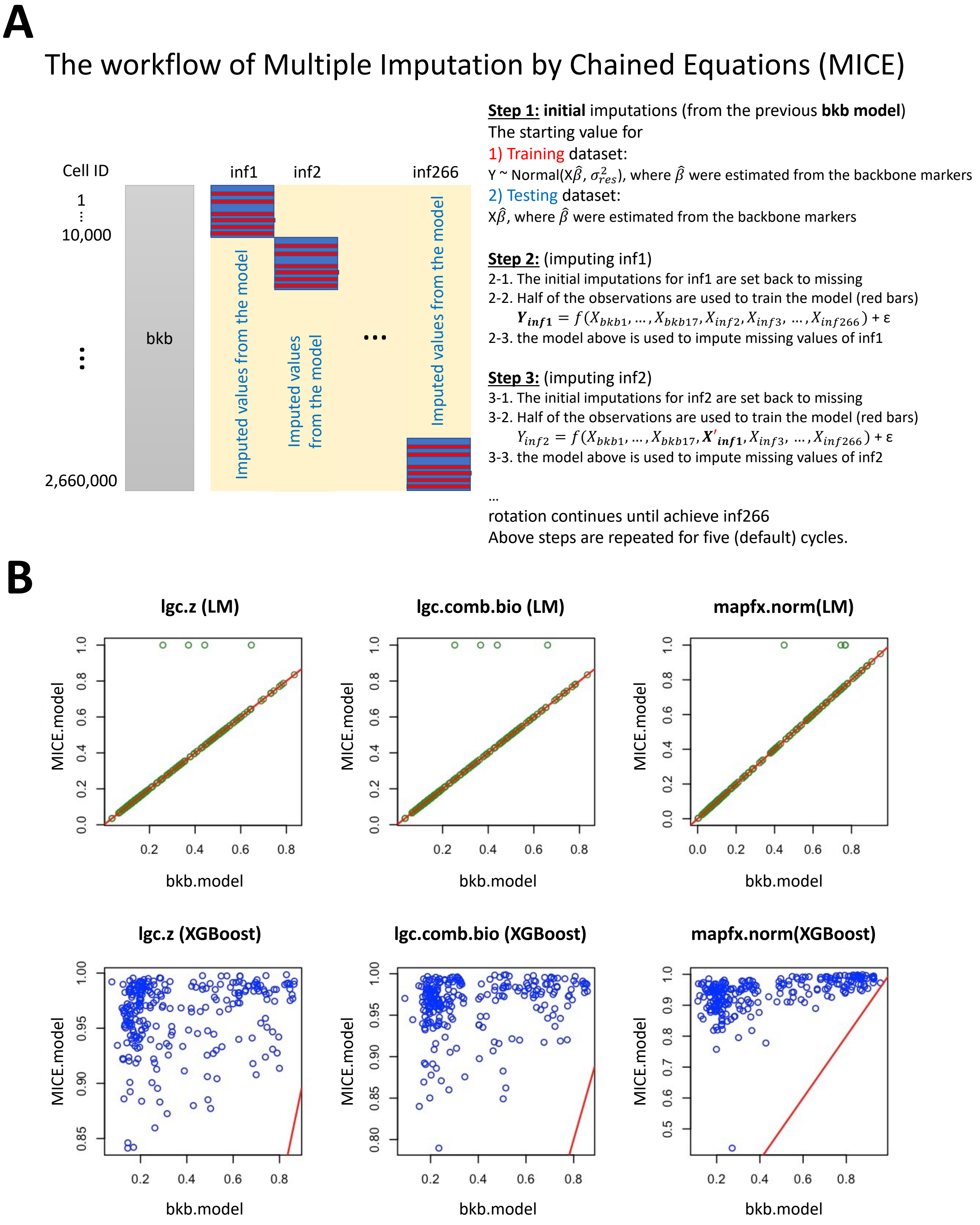


Figure 9: **The workflow of MICE and its model fit compared to imputations from the models with backbone markers as the only predictors.** (A) The workflow of MICE for the Becht *et al.* (2021) data. Each infinity marker gets updated by using both the *backbone* and the *infinity* markers as predictors with the starting values from the backbone model. The cells in the training set are highlighted in red, and the cells in the testing set are highlighted in blue in each diagonal block. (B) The $R^2$ values from the backbone model (x-axis) and the ones from the MICE framework (y-axis). Dark yellow dots in the upper row represent results from the linear model, and the blue dots in the lower row are the results from the XGBoost model. MICE improved the imputation greatly with the non-linear regression model (XGBoost), whereas there's no much improvement when using the linear regression model (LM).
